# Sec and Tat mediated secretion safeguards *Mycobacterium tuberculosis* membrane homeostasis

**DOI:** 10.1101/2025.09.04.674216

**Authors:** Priyadarshini Sanyal, Jagadeeshwari Uppada, Shashank Sinha, Yashasvi Bhat, Sidra Khan, Shri Vishalini Rajaram, Evanjalee Albert Arokiyaraj, Gagan Deep Jhingan, Nisheeth Agarwal, Areejit Samal, Vinay Kumar Nandicoori

## Abstract

Protein secretion drives *Mycobacterium tuberculosis* (*Mtb*) physiology and pathogenesis, yet a unified picture of the machinery and its role in cell membrane homeostasis is still lacking. By comprehensively curating published evidence, we assembled a systems-level map of *Mtb* secretion encompassing 92 components and 198 mechanistic reactions across Sec, Tat, and ESX pathways. The secretory components identified were integrated with high-throughput ChIP-Seq and transcriptome datasets to elucidate the regulation of the secretion system. Using CRISPRi, conditional depletion of SecA1 or TatA impaired growth *in vitro* and survival *ex vivo*. Quantitative secretome revealed decreased export of SecA1- and TatA-dependent substrates, with enrichment of cytosolic proteins in culture filtrates, indicating increased membrane permeability. Membrane proteomics showed depletion dependent increased metabolic/lipid-degrading proteins and decreased cell-wall/cell-process proteins, consistent with loss of membrane stability. Ultrastructural defects and increased ethidium bromide uptake confirmed impaired membrane integrity. Together, our multi-omics and functional genetics established SecA1 and TatA as essential guardians of *Mtb* membrane integrity which provided valuable datasets and a framework for secretion-dependent *Mtb* pathogenesis.

## Introductions

Protein secretion, a ubiquitous mechanism for transporting macromolecules across membranes, is found in all three domains of life: *Prokaryota, Archaea, and Eukaryota*. Eukaryotic protein secretion occurs through a general pathway called “Sec pathway,” which is present in the endoplasmic reticulum (ER) membrane and involves secretory vesicles carrying proteins to the Golgi apparatus^1^. In prokaryotes, the proteins are mainly secreted through two pathways, namely the Sec-dependent pathway^2,3^, and the Sec-independent, i.e., the twin-arginine translocation pathway (Tat pathway)^4^, which is also conserved in plants^5^. Sec or Tat pathways primarily translocate proteins from the cytoplasmic space to the periplasmic space, cell envelope, or extracellular milieu^6,7^. Proteins that are secreted through Sec and Tat pathways are identified by their respective N-terminal signal peptides^8^.

Sec-dependent secretion is divided into two types: a) co-translational translocation (Signal Recognition Particle (SRP)-mediated), which exports unfolded proteins while being translated by ribosomes, and b) post-translational translocation (SecA1 and SecA2-mediated), which exports unfolded proteins post-translation^9^. The Sec secretion system comprises SecY, SecE, and SecG proteins^9–11^, which form the SecYEG-SecDF-YajC-YidC holotranslocon, with the help of auxiliary proteins SecD, SecF, YajC, YidC^12^. SRP and SecA recognize the emerging preprotein with an N-terminal Sec-signal peptide. For SRP mediated transport, FtsY acts as a receptor and translocation occurs using GTP hydrolysis, whereas SecA as an ATPase, supplies energy for SecA mediated protein translocation^13^. Type I and II signal peptidases, associated with the holotranslocon, act on the cleavage sites, facilitating the conversion to a mature protein^8,14^. The twin-arginine translocation (Tat)^15^ system in prokaryotes and the thylakoid membrane of plant chloroplasts^4^ is involved in secreting a subset of folded proteins^15,16^ containing two invariant arginine residues (RR motif)^17^. In *E. coli*, five *tat* genes, i.e., *tatABCDE,* have been reported, among them *tatABCD* are present in an operon^18^.

*Mycobacterium tuberculosis* (*Mtb*) secretes an array of proteins, including virulence factors, with the help of generalized and specialized secretion pathways. In *Mtb*, the canonical SecA has two paralogs, SecA1 and SecA2^19,20^. While both SecA paralogs have ATPase activity, the non-canonical SecA2 differs from SecA1 in that it only secretes a small set of proteins to the extracellular space, which are known to be involved in virulence^21–24^. The *Mtb* genome does not encode *tatE*, and *tatD* is not involved in Tat-mediated protein export^18^. TatB and TatC form a Tat Complex-I (TC-I) in the inner membrane, and during translocation, bind to TatA to form Tat Complex-II (TC-II), which results in the formation of an oligomeric pore across the membrane that assists in protein translocation in the presence of proton motive force (PMF)^25^. Specialized secretion systems, such as the ESX secretion systems also known as Type VII Secretion systems (T7SS), transport small proteins with PE/PPE, WXG or YXXXD/E motifs^26^. T7SS consists of 5 systems, ESX-1 to ESX-5, which play an essential role in virulence, pathogenesis, iron, zinc homeostasis, and modulation of host-cellular processes^27^. Although each system has its own specific substrates and functions, they share essential structural features and a common evolutionary origin^28^. The widely studied ESX-1 system works in conjunction with ESX-3 and ESX-5. ESX-2 and ESX-4 are less characterized and are known to play important roles in DNA regulation^28^ and extracellular heme acquisition^29^.

There are several gaps in our understanding of this complex network of secretion systems in *Mtb* that we aimed to address. We asked: (i) what are the core and accessory components in different secretion systems, and how are they organized? (ii) How are these components involved in and regulated at multiple stages of protein transport? (iii) What is the role of SecA1 and Tat secretion systems in cell growth and survival, and what are their substrates? (iv) What is the impact of depleting SecA1 and TatA on *Mtb* membrane homeostasis? In this study, we employed a reconstruction approach to integrate molecular components with mechanistic interactions and developed a visual framework encapsulating the secretion process at the systems level. We generated conditional depletion strains of SecA1 and TatAC to explore their biological roles. Proteome analysis of wild-type and depleted strains suggested that, although the secretion of target substrates is compromised, it led to unusual secretion of cytosolic proteins. Experiments suggest that depletion of these systems impacts membrane homeostasis, reflected in changes in morphology and permeability.

## Results

### Reconstruction of Protein Secretion Systems in *Mtb*

The reconstruction of the secretory systems in *Mtb* was conducted as depicted in Fig. 1a-b. This comprehensive analysis inspired by the work Feizi *et al*. on genome-scale reconstruction of protein secretion system in yeast *Saccharomyces cerevisiae* aimed to elucidate context-specific regulation and pathway activity, providing insights into the secretion system’s behaviour across different physiological states. First, we performed an extensive and systematic literature review that encompassed research articles, review papers, and relevant book chapters on bacterial secretory pathways, with a focus on *Mtb*-specific literature. The evidence from the literature was further curated through a systematic query in the MycoBrowser database^30^. The final annotation is based on experimental evidence from either *Mtb* or other bacterial systems, computational predictions that include sequence similarity and domain architecture, and functional studies. This process resulted in a curated list of 92 key secretory components belonging to three different subsystems: Sec dependent (SRP mediated, SecA2 and SecA1 mediated), Sec independent (Tat), and T7SS (Table S1a). Among these, 19 components were unique to Sec pathways, 3 to Tat, and 67 to T7SS, respectively. Four components, namely LepB (Rv2903c), Lgt (Rv1614), LspA (Rv1539), and Ppm1 (Rv2051c), were shared between the Sec and Tat pathways and are primarily involved in the lipidation process and signal peptide cleavage. In contrast, five components, which are primarily chaperones (Rv3417c-GroEL1, Rv0440-GroEL2, Rv3418c-GroES, Rv0009-PpiA, and Rv2582-PpiB), were shared between the Tat and T7SS pathways (Fig. 1b). The structural and functional annotations of the components, including essentiality, subcellular localization, structural features, and biological functions, are listed in Table S1b, along with relevant evidences. The *in silico* predictions of the H37Rv proteome for secretory specific signal sequences resulted in 8 different categories of proteins belonging to Sec and Tat pathways (Fig. 1c). The proteins were categorized into different pathways depending on the presence of Sec/Tat specific signal peptide, lipo motif (LM) and transmembrane domain (TMD).

**Figure 1:**
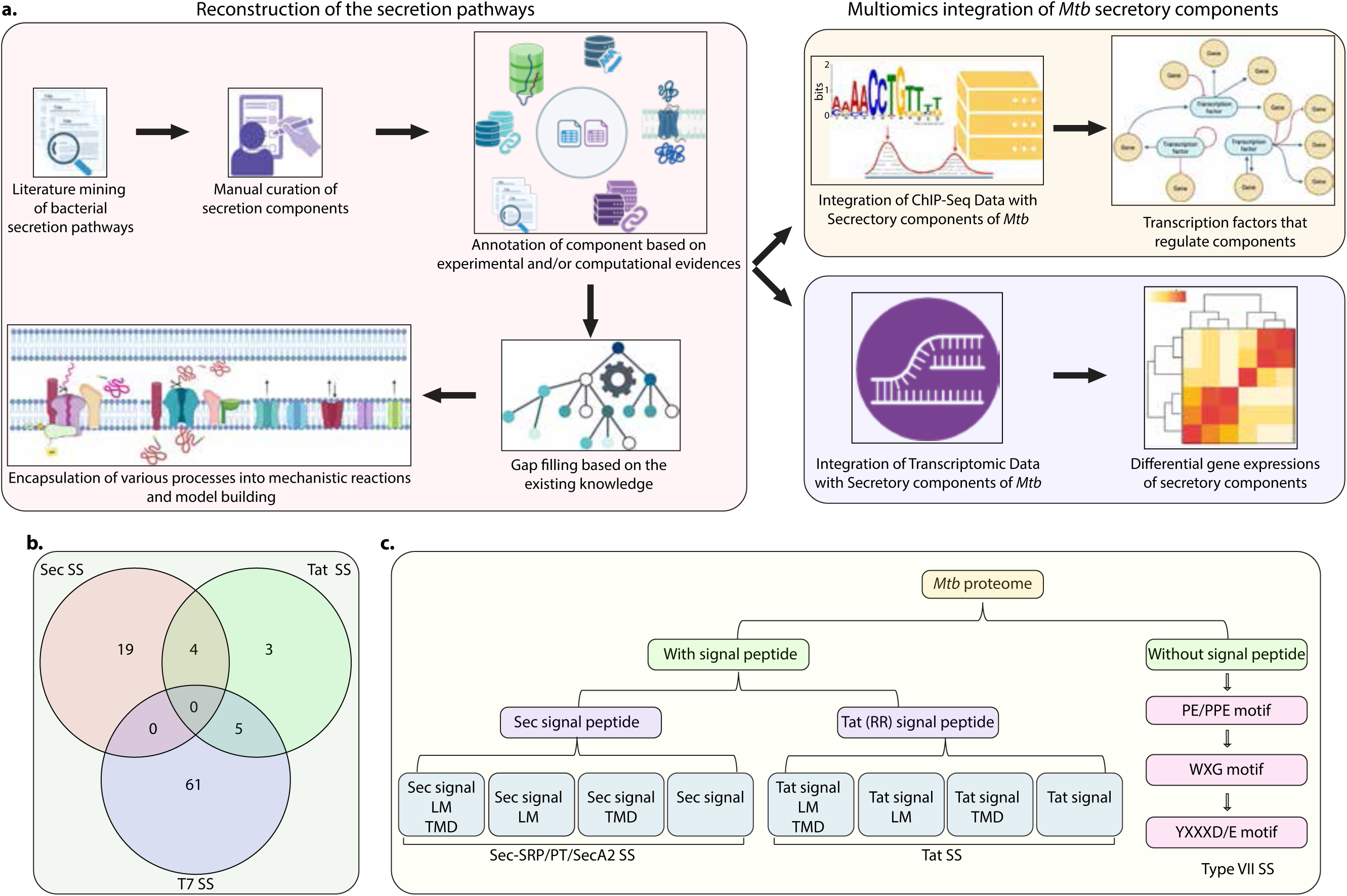
Reconstruction of Secretion pathways in *Mtb*. a) Workflow showing the steps involved in identification and annotation of components associated with secretion systems in *Mtb.* The secretory components were curated manually based on the experimental evidence and literature obtained from public sources. The integration of publicly available ChIP-Seq and RNA-Seq data with the curated list of components led to insights on regulation of secretion systems. b) Venn diagram depicting the distribution of the annotated components among the generalized secretion systems (Sec and Tat) and specialized (ESX1-5) secretion systems. c) *In silico* identification and characterization of *Mtb* proteins based on presence or absence of N-terminal signal sequence specific to secretory system, transmembrane domain (TMD) and Lipo motif (LM). The signal peptides and Lipo motifs specific to Sec and Tat were predicted by using <u>SignalP v6</u>, motifs specific to ESX system using <u>SecretoMyc</u> database, and transmembrane domains using <u>DeepTMHMM 1.0</u> and <u>Phobius</u>. After prediction of signatures, the proteins were divided into the categories depicted in the flowchart. Created in BioRender. Nandicoori, V. (2025) https://BioRender.com/n39ydxs.

The export mechanism for each protein category, based on the predictions and associated secretory components, was used to frame 198 mechanistic reactions that significantly represent the protein export process in *Mtb* (Table S2a and S2b). These 198 mechanistic reactions illustrate the step-by-step procedure involved in protein export, which was utilized to generate a predictive model of secretion systems in *Mtb* (Fig. 2 & Fig. S1-2). The three significant steps involved in protein export are the recognition of the N-terminal signal peptide in the nascent protein, the export of the protein through the respective translocon based on the signal sequences, and finally, the cleavage of the signal peptide, converting it into the mature protein. During translation, a nascent polypeptide with a hydrophobic signal sequence emerges from the ribosome. The SRP complex (Ffh bound to 4.5S RNA) recognizes and binds to the signal sequence and the ribosome-nascent chain complex (RNC). This binding temporarily halts translation to prevent improper folding. The SRP-RNC complex is then targeted to the membrane-bound SRP receptor, FtsY. Upon GTP binding and hydrolysis by Ffh and FtsY, the complex is transferred to the SecYEG translocon embedded in the plasma membrane. Translation resumes, and the growing polypeptide is inserted into or translocated across the membrane. Finally, SRP and FtsY dissociate and are recycled for subsequent targeting events (Fig. S2a-d).

**Figure 2:**
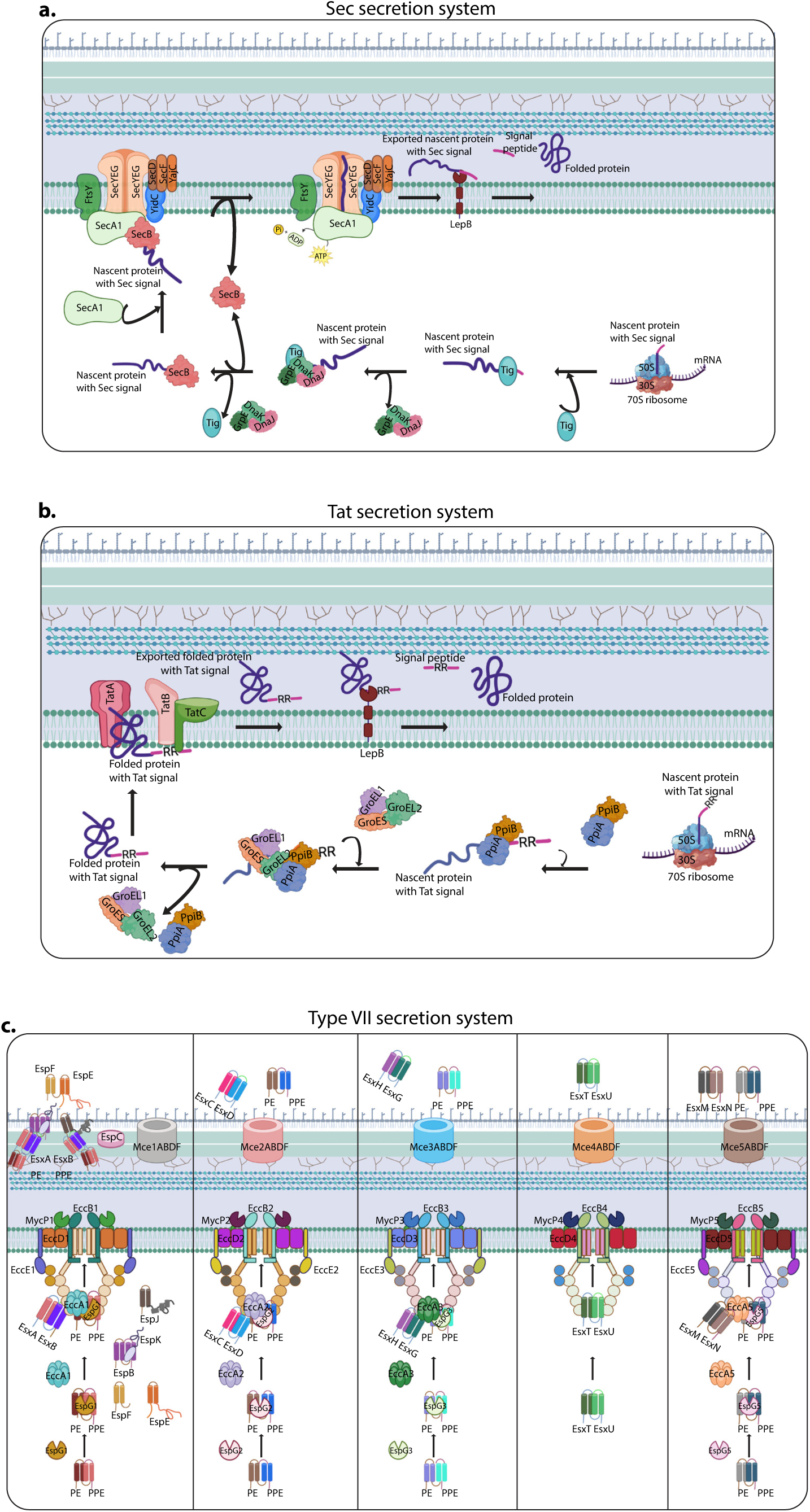
Schematic representation of secretion systems in *Mtb*. (a) Pictorial model depicting the steps involved in secretion of a protein containing N-terminal Sec specific signal peptide using SecA1 dependent Sec pathway. (b) Steps involved in secretion of a protein containing Tat signal peptide (twin arginine -RR- motifs) using Tat pathway. (c) Overview of protein secretion using specialized secretion systems (ESX1-ESX5) in *Mtb*. The steps shown were based on the mechanistic reactions framed using the experimental evidence provided in Table S2. Created in BioRender. Nandicoori, V. (2025) https://BioRender.com/oa5201s.

If a protein lacks an SRP-recognizable signal sequence, it typically follows alternative post-translational pathways for membrane targeting or secretion. One primary route is the SecA1-dependent pathway, where the fully synthesized protein, bound to the trigger factor (ribosomal chaperone), is recognized in the cytoplasm by SecA1, often with the help of chaperones like DnaK/DnaJ or SecB-like proteins that maintain the protein in an unfolded state. SecA1 then delivers the protein to the SecYEG translocon in the membrane and uses ATP hydrolysis to translocate it across (Fig. 2a). Alternatively, proteins with a twin-arginine motif are transported in a folded state via the TAT (Twin-Arginine Translocation) pathway (Fig. 2b). The twin-arginine motif (RR-motif) is recognized by TatC present in the complex with TatB and the translocation happens from the pore formed by TatA oligomerization in the inner membrane with the help of proton motive force.

The fate and localization of proteins depend on the presence of TMD or LM. If a protein contains a TMD, it is embedded into the inner membrane using the membrane insertase (YidC) during translocation (Fig. S1). If the protein does not have an LM, the signal peptidase LepB cleaves the N-terminal signal peptide. For proteins containing a LM, additional steps of lipidation occur in the inner membrane, involving three enzymes: Lgt (Phosphatidylglycerol: prolipoprotein diacylglyceryl transferase), which adds a diacylglycerol to the thiol of the lipobox cysteine located after the signal sequence; LspA (Signal peptidase II), which cleaves the signal peptide, exposing the lipid-modified cysteine at the N-terminus; and Ppm1, which adds a third amide-linked fatty acid, completing triacylation. The mature triacylated lipoprotein is correctly anchored and functional at the outer membrane or retained in the inner membrane, depending on the transmembrane domains (Fig. S1).

In the case of *Mtb*, a smaller subset of proteins that lack classical signal peptides or have atypical ones, such as SodA (superoxide dismutase)^21^, KatG (catalase-peroxidase)^21^, SapM (Acid phosphatase)^22^, PknG (Serine/threonine-protein kinase-G)^31^, and certain PE/PPE proteins, are transported through SecA2 transport. SecA2 is predominantly cytosolic and exports proteins in conjunction with the chaperone SatS^32^. SecA2 may function by interacting with SecYEG either independently or in cooperation with SecA1^33^. Some evidence suggests that SecA2 may require SecA1 for effective translocation, possibly forming heterodimers or coordinating the sequential export of substrates^34^. SecA2 reactions were not integrated into the model, as there is no clear evidence of how these proteins are transported. Unlike the canonical Sec system, SecA2 substrates often fold fully or partially before export^33,35^.

In addition to the aforementioned secretion systems, *Mtb* also contains ESX-1 to ESX-5^36^ secretion systems that facilitate the transport of virulent proteins across the membrane to an extracellular environment^37,38^. The proteins transported through ESX are known to feature either a PE/PPE motif or a WXG motif at the N-terminal and a YXXXD/E motif at the C-terminal^26^. Studies have demonstrated that components of the ESX systems create a channel in the inner membrane (Fig. 2c), although the mechanism by which the proteins are exported into the extracellular environment remains unclear. However, recent proteogenetic studies conducted by Cronin et al., 2022^39^ showed that the substrates of ESX-1 are co-secreted, and the absence of one gene leads to the loss of both secreted complexes, also outlining the order in which these substrates are transported as described in Table S2.

### Integration of Omics data with Secretory Components

The transcriptional regulation of secretory components was analyzed by curating and integrating publicly available ChIP-Seq data^40,41^ along with the secretory components listed in this study. This integrative analysis revealed that 68 Transcription Factors (TFs) from 28 families were directly involved in regulating secretory components. Most of the TFs belonged to the TetR family, followed by the ArsR family. Among the TFs, 50% (34 of 68) were known to regulate more than four secretory components (Fig. 3a). The major TFs involved in this regulation include Rv0081, which belongs to the ArsR family and directly regulates 12 components within the Sec pathway (6), Tat pathway (1), and T7SS (5). The second TF that controls most secretory components is *cosR* (*rv0967*), which plays a vital role in copper homeostasis and directly regulates 11 secretory components. Lastly, *trcR* (*rv1033c*), a response regulator of the two-component system TrcRS, directly regulates nine secretion components (Fig. 3b).

**Figure 3:**
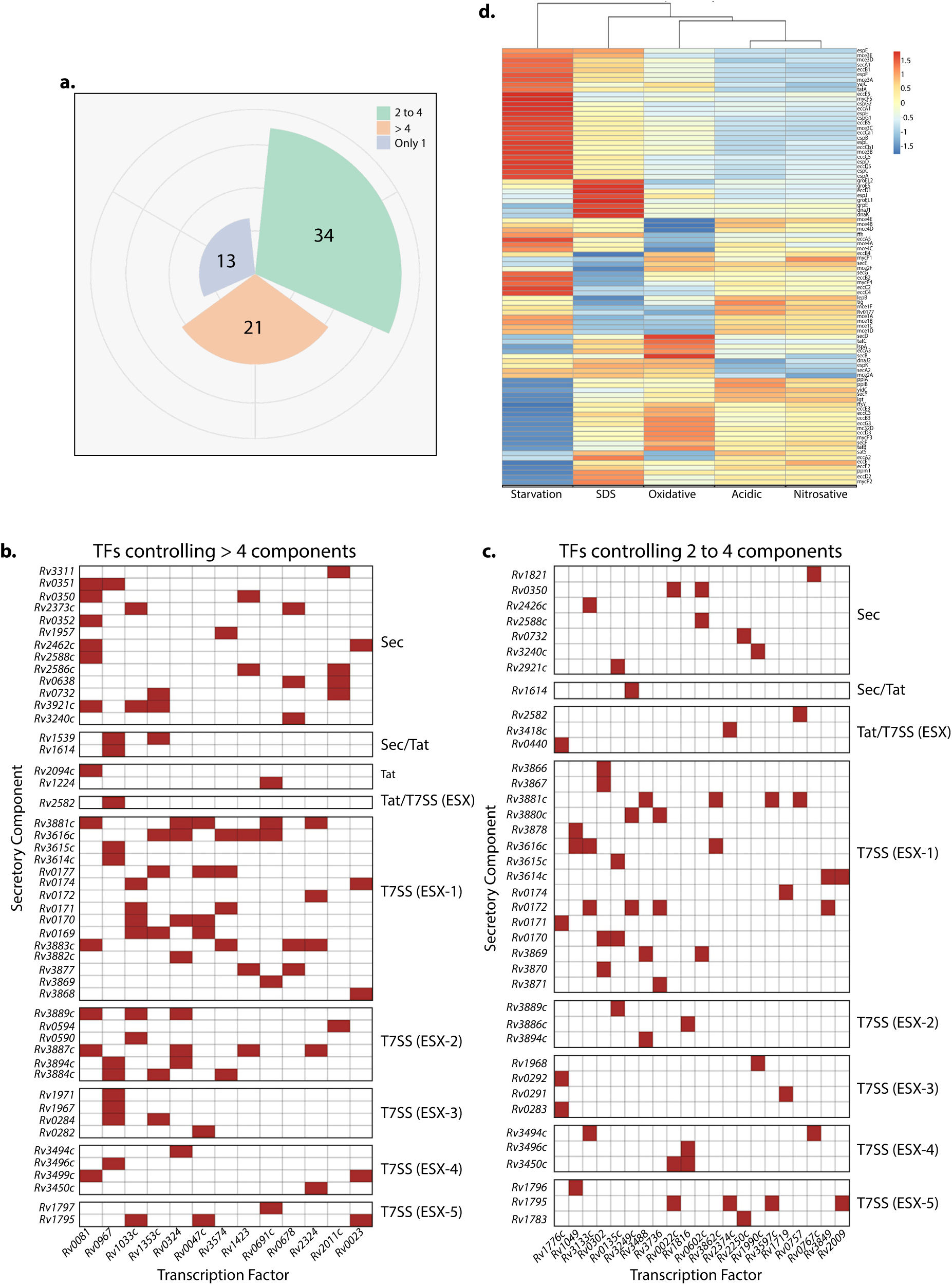
Transcriptional regulation of components of secretion systems and differential expression of secretory components under different stress conditions. (a) Radial pie chart grouping the transcription factors (TFs) based on the number of the secretory components they directly regulate. Checkerboard plots illustrating the regulation of secretory components by transcription factors. (b) TFs controlling > 4 components and (c) TFs controlling 2-4 components. Each row represents a Transcription Factor (x-axis) and each column a regulated secretory component (y-axis). The secretory components were grouped based on the three secretion systems. Common secretory components shared by the secretion systems were also shown. Presence of a filled square indicates a detected direct regulatory interaction. The ChIP-Seq data for TFs was obtained from <u>MTB Network portal.</u> (d) Heatmap depicting the differential expression (Log2FC) of the secretory components under different stress conditions. The differential expression values were obtained from RNA Seq data published by <u>Khan et al., 2024</u>.

Among the 21 factors regulating the expression of more than 2 secretory components and fewer than four components, most belong to the TetR and WhiB families and are primarily observed in controlling the T7SS pathway (Fig. 3c). The expression of secretory components is mainly associated with external stimuli. The stress conditions faced by the organisms aid in secreting various proteins. Therefore, we aimed to examine the transcriptional level expression of the secretory components under several stress conditions, including oxidative, nitrosative, SDS, and acidic stress. This can be achieved by integrating the RNA sequencing data^42^ with the secretory components data. We looked at the differential expression of secretory components under various stress conditions compared to their expression in cells grown in 7H9 medium (Control). We observed that starvation and SDS stress significantly impacted the expression of these secretory components relative to other stresses. Most of the Sec and Tat components were downregulated under starvation stress, whereas the opposite was true for the T7SS (Fig. 3d).

### Compromised bacterial survival upon *secA1* and *tatAC* depletion

T7SS systems in *Mtb* are extensively investigated in relation to other secretion pathways, specifically the Sec and Tat pathways. The deletion of SecA2 and its impact on the secretome has already been reported. Therefore, in this report, we aim to focus on and investigate the SecA1 and Tat secretion pathways; thus, we selected SecA1 and TatA as the candidates. Furthermore, both SecA1 and TatA are known to be regulated by PknB-mediated phosphorylation^43^, which makes them intriguing candidates. High-throughput transposon mutagenesis studies indicate that both *secA1* and *tatA* are essential genes for *in vitro* growth in complete media.

We generated CRISPRi-mediated knockdown strains^44^ of *secA1* and *tatA*. Towards this, either the pTetIntdCas9 vector or the pTetIntdCas9 harboring two guide RNAs (Fig. S3a) targeting *secA1* or *tatA* (Fig. S3b) were electroporated into *Mtb* to create *Rv-vc_dc_, Rv-secA1_dc_*, and *Rv-tatA_dc_* strains. The addition of ATc results in the expression of dCas9 and the guide RNA, along with their handle, which form a complex and stall transcription (Fig. S3c). The depletion of SecA1 and TatA upon the addition of ATc and the expression of the were confirmed by Western blotting with α-SecA1 and α-TatA antibodies, respectively (Fig. 4b). Depletion of either SecA1 or TatA resulted in compromised growth *in vitro* (Fig. 4a).

**Figure 4:**
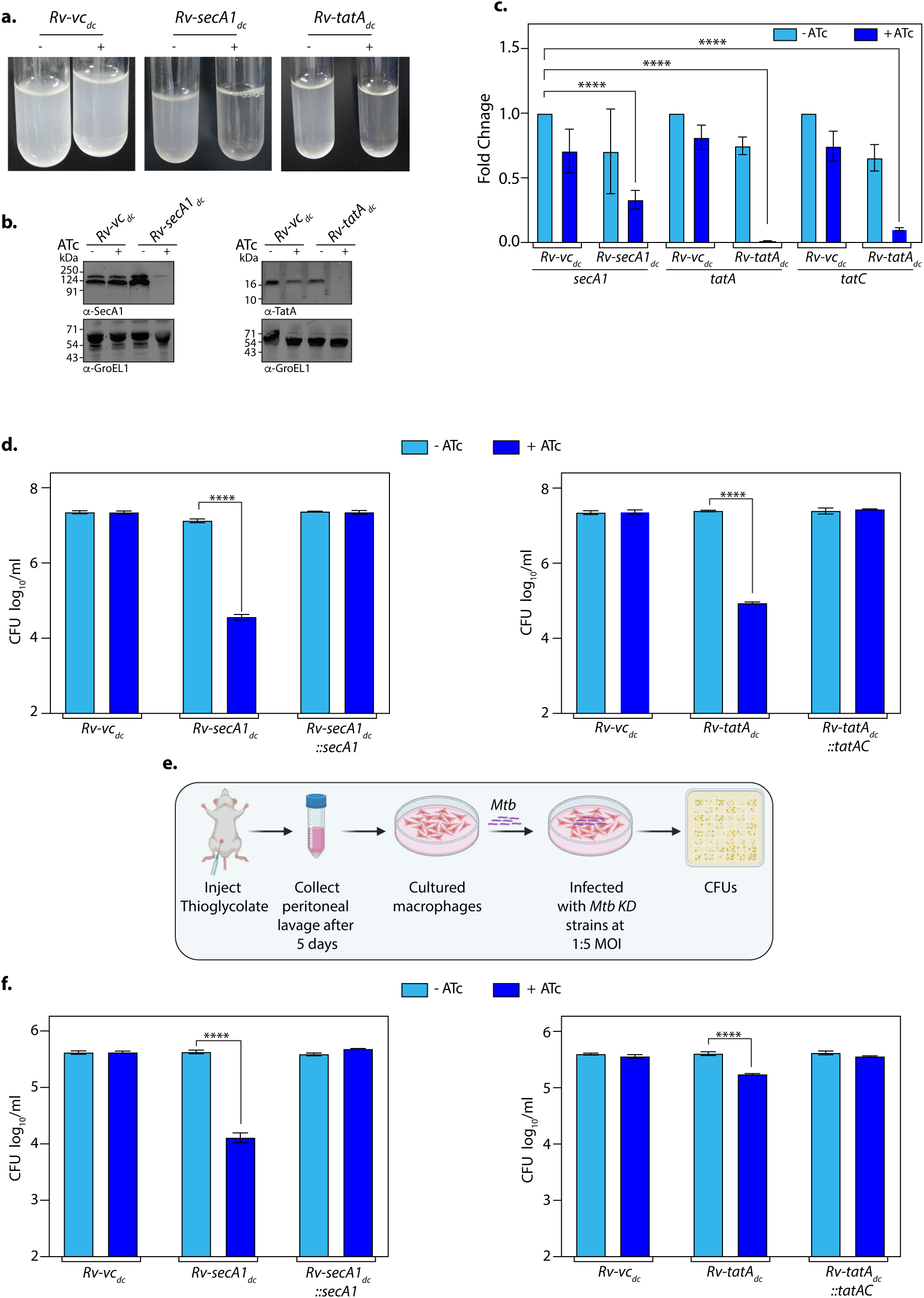
Impact of bacterial survival upon *secA1* and *tatA* depletion. a) *Rv-vc_dc_*, *Rv-secA1_dc_* and *Rv-tatA_dc_* were inoculated at A_600_ ∼0.1 and grown for 4 days in the presence and absence of ATc. The tubes represent the depletion of *Rv-secA1_dc_* and *Rv-tatA_dc_* as compared to *Rv-vc_dc_*. b) Whole cell lysates (WCLs) were prepared from *Rv-vc_dc_*+ATc, *Rv-secA1_dc_*-/+ATc and *Rv-tatA_dc_*-/+ATc from grown 4^th^ day culture. 50 µg of each protein lysates were resolved on 10% (SecA1) 15% (TatA) SDS-PAGE, transferred on Nitrocellulose membrane and probed with α-SecA1 (for SecA1 depletion), and α-TatA (for TatA depletion) antibodies generated in the lab. α-GroEL1 antibody was used as loading control. c) Total RNA was isolated from *Rv-vc_dc_* -/+ATc, *Rv-secA1_dc_* -/+ATc and *Rv-tatA_dc_* -/+ATc from 3^rd^ day grown cultures for qRT-PCR. The expression of *secA1, tatA* and *tatC* were normalized with respect to 16S rRNA and the Fold Change (2^-ΔΔCt^) value (mean ± SD (n = 4)) was plotted using GraphPad Prism. The Statistical significant was analysed using One-way ANOVA, ∗∗∗, p < 0.0001. d) The graph depicts the CFU of the *in vitro* survival of bacteria of *Rv-vc_dc_* -/+ATc, *Rv-secA1_dc_*-/+ATc, *Rv-secA1_dc_::secA1* -/+ATc, *Rv-tatA_dc_*-/+ATc and *Rv-tatA_dc_::tatAC* -/+ATc on 4^th^ day post depletion. GraphPad Prism software was used to plot CFUs and performed statistical analysis using One way ANOVA. Three independent experiments were performed as biological replicates and each experiment was conducted in triplicates. Data represents mean and ± SD, **** p<0.0001. e) Schematic representation of peritoneal macrophages isolation from mice peritoneum. 4 days post thioglycolate injection at the mice peritoneum, the macrophages were collected, seeded for 16h and infected with the *Rv-vc_dc_, Rv-secA1_dc_, Rv-secA1_dc_::secA1, Rv-tatAdc,* and *Rv-tatAdc::tatAC* at 1:5 MOI. 4 h post infection, CFUs were enumerated at 0h and 96h. Created in BioRender. Nandicoori, V. (2025) https://BioRender.com/xwcvl0i. f) Survival of the *Rv-vc_dc_, Rv-secA1_dc_, Rv-secA1_dc_::secA1, Rv-tatA_dc_*, and *Rv-tatA_dc_::tatAC* in the present and absence of ATc. GraphPad Prism software was used to plot CFUs and performed statistical analysis using One way ANOVA. Three independent experiments were performed as biological replicates and each experiment was conducted in triplicates. Data represents mean and ± SD, **** p<0.0001.

The CRISPRi system is known to cause polarity effects for the entire operon^44^. While *secA1* is a solo gene, *tatA* is the first gene in a two-gene operon that also includes *tatC*. Both *tatA* and *tatC* are essential *in vitro* and are part of the Tat secretion pathway. To evaluate the polarity effect, total RNA was isolated from the *Rv-vc_dc_, Rv-secA1_dc_*, and *Rv-tatA_dc_* strains in the presence and absence of ATc, and the expression levels of *secA1*, *tatA,* and *tatC* were quantified (Fig. 4c). Upon the addition of ATc, although we were targeting only *tatA*, the expression levels of both *tatA* and *tatC* were compromised. Thus, we used *tatA_gmut_* alongside *tatC* to complement *Rv-tatA_dc_*. We electroporated complementation constructs pSNG-secA1_gmut_ or pSNG-tatAC_gmut_, in which the guide RNA target sequences were mutated (Fig. S3d), into *Rv-secA1_dc_* and *Rv-tatA_dc_* strains, respectively, to generate *Rv-secA1_dc_::secA1* and *Rv-tatA_dc_::tatAC* (Fig. S3e).

The functional complementation was validated by enumerating CFU *in vitro* and in *ex vivo*. The cells were grown in 7H9 media, and CFUs were enumerated at different time points. In the *Rv-secA1dc* +ATc strain, compromised growth was observed around ∼4 log_10_ fold as compared to *Rv-vc_dc_* +ATc after 4 days of depletion (Fig. 4d). While in the case of *Rv-tatA_dc_* +ATc, the growth defect was noticed around ∼3 log_10_ fold (Fig. 4d). The growth phenotypes were rescued by complementation. In the peritoneal macrophages (Fig. 4e), the growth was compromised up to ∼2.5 log_10_ fold in *Rv-secA1_dc_* +ATc and ∼0.8 log_10_ fold in *Rv-tatA_dc_* +ATc as compared to *Rv-vc_dc_*+ATc control (Fig. 4f). The complementation strains rescued the growth phenotype. We conclude that we successfully generated the *Rv-secA1_dc_* and *Rv-tatA_dc_*knockdown strains, indicating that *secA1* and *tatAC* (*tatA* and *tatC*) are essential for bacterial survival.

### Change in the secretome upon SecA1 and TatAC depletion

SecA1 and TatAC are important players involved in Sec and Tat-mediated secretion pathways. While *in silico* studies provide predictions on putative Sec or Tat substrates based on the presence of consensus signal peptides or twin arginine motifs, respectively, there are no experimental studies that confirm these predictions. To identify the SecA1- and Tat-mediated secretome, *Rv-vc_dc_*, *Rv-secA1_dc,_* and *Rv-tatA_dc_* strains were grown to the early log phase (A_600_ ∼0.4) in the presence of ATc. Cells were spun down, the pellet was processed for WCL, and the supernatant was enriched for the culture filtrate (CF) proteins^45^ (Fig. S4a). The WCL and CF fractions were resolved and probed with α-PknB and α-Ag85B antibodies, which are markers for membrane/WCL and secreted proteins, respectively (Fig. S4b). Absence of band corresponding to PknB and presence of Ag85B bands in CF fractions is indicative of the lack of contamination from other fractions. Ag85B is shown to be secreted through SecA pathway in *E. coli*^46^. As anticipated, we observed that Ag85B levels were lower upon the depletion of SecA1; however, we also observed decreased levels of Ag85B upon TatAC depletion. We cannot rule out the possibility that Ag85B may use more than one secretory pathway for its secretion.

The CF fractions in triplicates were trypsinized, and the resulting tryptic peptides were resolved using LC-MS/MS, followed by data analysis with label-free quantitation (LFQ). The data was further processed as outlined in Fig. 5a. A total of 1255 proteins were present in all three samples namely: *Rv-vc_dc_* +ATc, *Rv-secA1_dc_* +ATc, and *Rv-tatA_dc_* +ATc. Proteins with ≤ 2 unique peptides or with more than one transmembrane helix were filtered out, which decreased the number to 1010 proteins. This was followed by filtering to remove proteins

**Figure 5:**
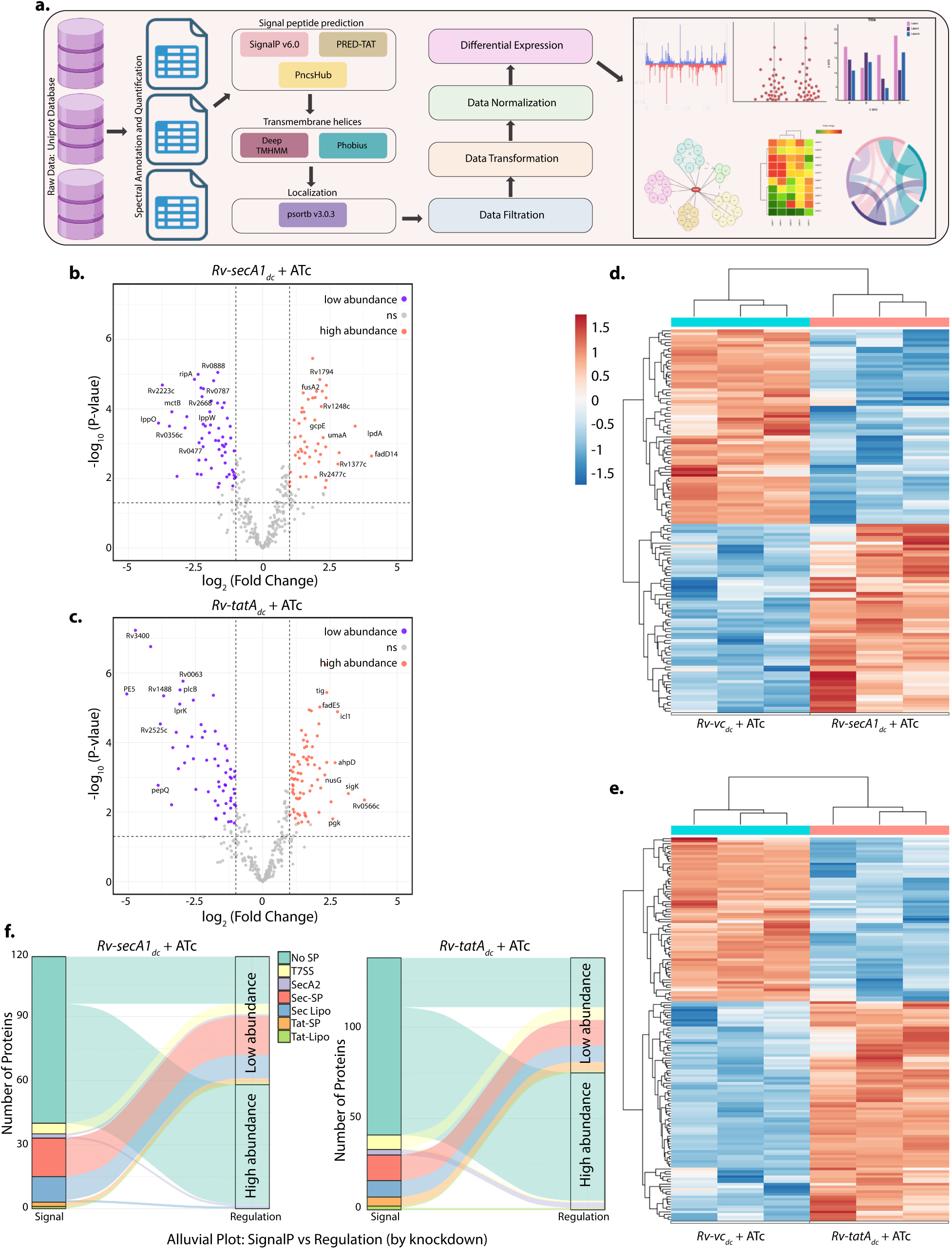
Secretome of SecA1 and TatAC. a) Schematic representation of Secretome data analysis of SecA1 and TatAC. Raw data was analysed using Uniprot database. The data was screened for spectral annotation. The proteins were predicted for signal peptides for Sec, Tat and T7SS using <u>SignalP v6</u>, PRED-TAT and PncsHub respectively, followed by removal of transmembrane helices using <u>DeepTMHMM</u> <u>1.0</u> and <u>Phobius</u>. Localization of the proteins were also checked using psortb v3.0.3. The filtered proteins were normalised and differential protein abundances were plotted. Created in BioRender. Nandicoori, V. (2025) https://BioRender.com/v0npynh. b-c) The volcano plots of SecA1 (b) and TatAC (c) depletion using 1≥log_2_Fold Change≤-1. The purple dots depict the low abundance proteins, orange dots depict the high abundance proteins and grey dots represent non-significant proteins. d-e) The heat maps depict the differential protein abundances in *Rv-vc_dc_*+ATc, *Rv-secA1_dc_*+ATc (d) and *Rv-vc_dc_*+ATc, *Rv-tatA_dc_*+ATc (e) using padj<0.05. The blue colour indicates the high abundant proteins and the orange colour represents the low abundance proteins. f) The Alluvial plots represent the number of proteins containing various signal peptides and their abundances upon SecA1 and TatAC depletion.

identified as part of membrane vesicles, which further decreased the number to 932 proteins. The resultant secretome obtained for *Rv-secA1_dc_* +ATc and *Rv-tatA_dc_* +ATc was compared with *Rv-vc_dc_* +ATC to identify the differences. We hypothesized that the absence of SecA1 or TatAC would lead to a substantial decrease in the number of secreted proteins. Contrary to our hypothesis, we observed a higher number of secreted proteins, which included known cytosolic proteins in both *Rv-secA1_dc_* +ATc and *Rv-tatA_dc_* +ATc samples, which could be due to changes in overall cellular homeostasis upon depletion (Table S5).

Therefore, to accurately analyze the changes in the secretome, we limited our analysis to secretory proteins that were common between the current study and at least one other previously published study (among the four previous studies^47–50^) (Fig. S4d). This reduced the number of secretory proteins selected for further analysis from 932 to 439. We compared *Rv-vc_dc_* +ATC with *Rv-secA1_dc_* +ATc or *Rv-vc_dc_* +ATC with *Rv-tatA_dc_* +ATc samples, and only those with a ±2 fold change were considered for further analysis. We obtained a total of 134 proteins that were differentially secreted in *Rv-secA1_dc_* +ATC, with 66 showing >2 fold higher abundance and 68 showing lower abundance (Fig. 5b). The numbers were 158 differentially secreted proteins, with 81 showing >2 fold higher abundance and 77 showing lower abundance for the *Rv-tatA_dc_* +ATc samples (Fig. 5c). Applying a p-value cut off <0.05 for the differentially abundant secretory proteins reduced the final counts to 118 (58 with higher and 60 proteins with lower abundances) for *Rv-secA1_dc_* +ATC and 138 (75 proteins with higher abundance and 63 with lower abundance) for *Rv-tatA_dc_* +ATc samples (Fig. 5d & e). Among these, 72 proteins were common between *Rv-secA1_dc_* +ATc and *Rv-tatA_dc_* +ATc samples, with proteins exhibiting similar increased or decreased abundance patterns in both samples (Fig. S4e).

We reasoned that the proteins less abundant in *Rv-secA1_dc_* +ATc but not in *Rv-tatA_dc_* +ATc samples are specific substrates of the SecA1 pathway (Fig. 4Sd-27 proteins), and those less abundant in *Rv-tatA_dc_*+ATc but not in *Rv-secA1_dc_*+ATc are specific Tat pathway substrates (Fig. 4Sd-30 proteins). The remaining 33 less abundant proteins (Fig. 4Sd-33 proteins) are either substrates of both pathways or require components from both pathways for efficient secretion. Among the 27 unique substrates of the SecA1 pathway, 13 proteins had SecA1 signal peptide (SP) and SP + LM (Fig. 5f). Among the 30 unique substrates of the Tat pathway, only 5 proteins were found to have a predicted Tat signal and Tat signal + LM. These results indicate that many of the less abundant proteins identified in the current study are not predicted to be substrates of these pathways, which may require further validation.

We also observed that a total of 58 and 75 secretory proteins were highly abundant (>2 fold and p-value <0.05) in *Rv-secA1_dc_* +ATc and *Rv-tatA_dc_* +ATc (Fig. 5d-e; Table S5). Among these, 39 proteins were common to both samples, while 19 were unique to *Rv-secA1_dc_* +ATc and 36 were unique to *Rv-tatA_dc_* +ATc. Of the 58 abundant proteins in *Rv-secA1_dc_* +ATc, 57 did not possess any N-terminal signal peptides. Proteins predicted to be secreted via T7SS and SecA2 pathways were highly abundant in *Rv-tatA_dc_* +ATc. We speculate that the higher abundance of proteins in *Rv-secA1_dc_* +ATc and *Rv-tatA_dc_* +ATc samples could be due to increased non-classical secretion (T7SS or SecA2). Furthermore, there is a possibility that depletion of the SecA1 and TatAC, known to be associated with the export of proteins involved in membrane stabilization, could have compromised the *Mtb* membrane integrity, leading to leakage of periplasmic or cytoplasmic proteins into the CF fraction. To address this, we investigated the changes in the membrane proteome upon SecA1 and TatAC depletion.

### Mycobacterial membrane proteome alteration owing to SecA1 and TatAC depletion

Next, we examined the protein composition of membrane fractions from *Rv-vc_dc_* +ATc, *Rv-secA1_dc_* +ATc, and *Rv-tatA_dc_* +ATc samples. The cytosolic and membrane fractions were probed with α-rpoA (a marker for the cytosolic fraction) and α-PknB (a marker for the membrane fraction) antibodies, respectively. The presence of PknB band and RpoA band in the WCL fraction, but not in cytosolic or membrane fractions, respectively, in *Rv-vc_dc_* +ATc samples suggests the purity of the respective fractions and validates the protocol used. While the fraction of *Rv-tatA_dc_*+ATc showed a profile similar to *Rv-vc_dc_* +ATc, we observed the presence of the cytosolic marker in *Rv-secA1_dc_* +ATc in the membrane fraction (Fig. S5b). The fractions were probed with other cytosolic markers, which supported the above observations (data not shown).

The membrane fractions in triplicate were trypsinized, and the samples were resolved by LC-MS/MS, with data processed through label-free quantitation (LFQ) and analyzed as described in Fig. 6a. A total of 2335 proteins were detected in the membrane fractions *of Rv-vc_dc_* +ATc, *Rv-secA1_dc_* +ATc, and *Rv-tatA_dc_* +ATc samples. Proteins with ≤2 unique peptides and lacking transmembrane helices were eliminated from 2335 proteins, reducing the number of proteins to 1101. We compared the data obtained from the current study with two previously published membrane proteome studies and identified 943 proteins that were common to two studies^22,51^ including our study. We analyzed the abundances of these proteins in *Rv-secA1_dc_* +ATc and *Rv-tatA_dc_* +ATc samples in comparison with *Rv-vc_dc_*+ATc, and the proteins with a ±2 fold change, and with a p-value cut off of <0.05 were considered for further analysis.

**Figure 6:**
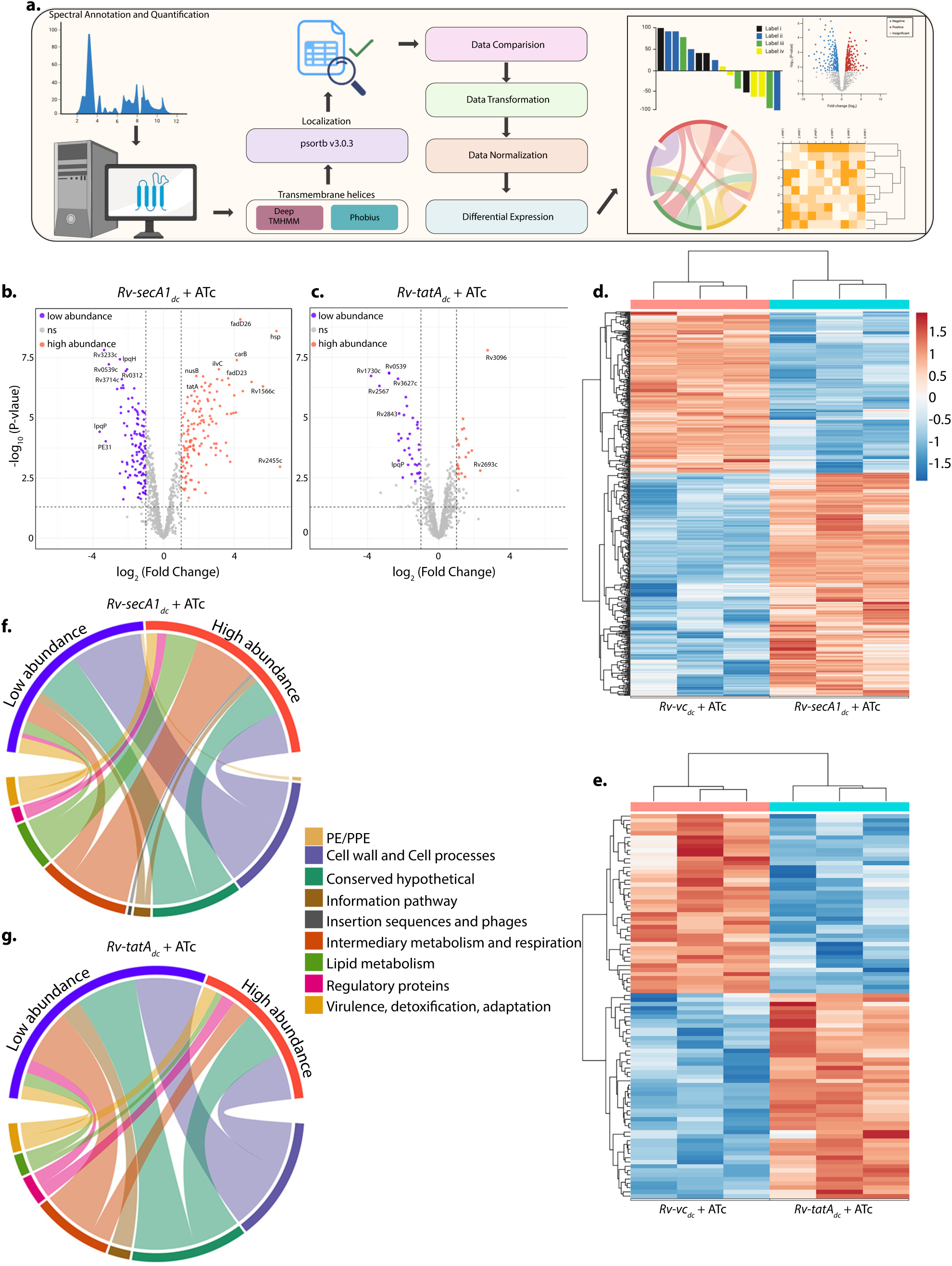
Membrane proteome of SecA1 and TatAC depleted cells. a) Schematic diagram of membrane proteome data analysis of SecA1 and TatAC. Raw data was analysed using Uniprot database. The data was screened for spectral annotation and quantification. The transmembrane proteins were selected using <u>DeepTMHMM 1.0</u> and <u>Phobius</u>. Localization of the proteins were also checked using psortb v3.0.3. The filtered proteins were normalised and differential protein abundances were plotted. Created in BioRender. Nandicoori, V. (2025) https://BioRender.com/u4nhpba. b-c) The volcano plots represent the proteins of SecA1 and TatAC depletion using padj<0.05 and 1≥log2Fold Change≤-1. The purple dots depict the low abundance proteins, orange dots depict the high abundance proteins and grey dots represent non-significant proteins. d-e) The heat maps depict the differential protein abundances in *Rv-vc_dc_*+ATc, *Rv-secA1_dc_*+ATc (d) and *Rv-vc_dc_*+ATc, *Rv-tatA_dc_*+ATc (e) membrane proteins. The blue colour depicts the high abundant proteins and the orange colour represents the low abundance proteins. f) The chord plot depicts the abundances of high and low proteins upon SecA1 depleted condition according to the different functional categories. g) The chord plot depicts the abundances of high and low proteins upon TatAC depleted condition according to the different functional categories.

We observed a total of 260 proteins that were differentially abundant in *Rv-secA1_dc_* +ATc, with 138 and 122 proteins showing high and low abundance, respectively (Fig. 6b & d). The number of differentially abundant proteins identified in *Rv-tatA_dc_* +ATc was much lower, accounting for 81 in total, with 31 being highly abundant and 50 low in abundance (Fig. 6c & e). The results seem to reflect the data observed in western blot (Fig. S5b), where cytosolic markers were detected in *Rv-secA1_dc_* +ATc samples, unlike in *Rv-tatA_dc_*+ATc. A total of 32 differentially abundant proteins were common in *Rv-secA1_dc_*+ATc and *Rv-tatA_dc_*+ATc samples. In *Rv-secA1_dc_* +ATc, 19 were high and 13 were low in abundance, while, 15 proteins were high and 17 were low abundant in case of *Rv-tatA_dc_* + ATc (Table 6). Among these 32 proteins, Phenolpthiocerol synthesis type-I polyketide synthase (PpsE), Proteasome beta subunit (PrcB), Rv0455c, and Rv1896c, were highly abundant in *Rv-secA1*_dc_ +ATc, whereas they were low in *Rv-tatA*_dc_ +ATc (Table 6). Annotation of differentially abundant proteins based on their functions showed that proteins belonging to cell wall and cell processes were low in abundance in the membrane fraction, while those involved in metabolic processes were more abundant in *Rv-secA1_dc_* +ATc (Fig. 6f). However, both cell wall and cell process proteins, along with metabolic process proteins, were low in abundance in *Rv-tatA_dc_* +ATc (Fig. 6g). Together, the data suggests that there may be a possible membrane dysbiosis in *Rv-secA1_dc_* +ATc samples, which may also be the case with *Rv-tatA_dc_* +ATc, but at a lower scale.

### SecA1 and TatAC depletion results in Mycobacterial membrane permeability

Data presented in Fig. 5 and 6 showed significant changes in the secretome and membrane proteome upon SecA1 and TatAC depletion. Interestingly, we observed many cytosolic proteins in both the secretome and membrane proteome when compared with Rv. Furthermore, we also observed that many proteins were common to both pathways, which are thought to be distinct. Thus, we speculated that the depletion of SecA1 to a large extent and TatAC to a lesser extent may be compromising the membrane, which could have led to the similarity. To investigate the phenotypic impact of SecA1 and TatAC depletion on cellular morphology, we performed scanning electron microscopy (SEM). We observed bulged and elongated cells (>40% cells were >3μm ) upon depletion of SecA1, which could be restored through complementation (Fig. 7a-b & Fig. S6a). While the cells were elongated (>10% cells were >4μm ) compared with Rv upon TatAC depletion, which could be complemented, they did not exhibit the bulged phenotype observed upon SecA1 depletion (Fig. 7a-b & Fig. S6a-b). To further investigate the effect of SecA1 and TatAC depletion on the ultrastructure of the *Mtb* membrane, we performed transmission electron microscopy (TEM). We observed that the architecture of the cell membrane was significantly altered upon SecA1 or TatAC depletion and could be restored upon complementation (Fig. 7c). However, we could not observe perforations in the membrane, suggesting that the possible changes at the given time point in the experiment may be more subtle and possibly more permeable.

**Figure 7:**
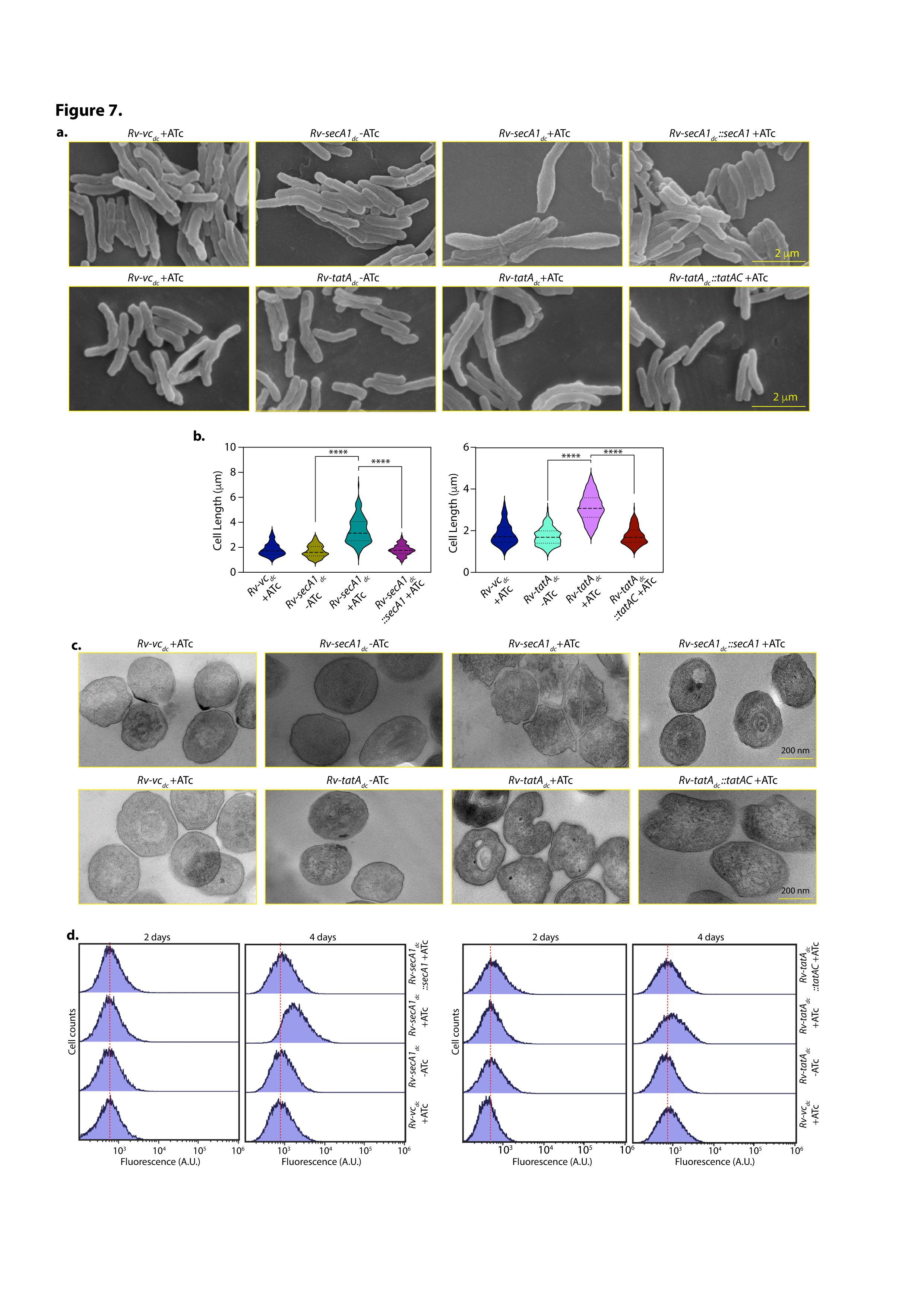
Alteration of the cellular morphology upon SecA1 and TatA depletion. a) SEM images of *Rv-vc_dc_* +ATc*, Rv-secA1_dc_* -/+ATc*, Rv-secA1_dc_::secA1* +ATc*, Rv-tatA_dc_* -/+ATc, and *Rv-tatA_dc_::tatAC* +ATc*. Rv-vc_dc_* +ATc*, Rv-secA1_dc_* -/+ATc*, Rv-secA1_dc_::secA1* +ATc*, Rv-tatA_dc_* -/+ATc*, and Rv-tatA_dc_::tatAC* +ATc were grown for 4 days post ATc addition, cells were fixed and processed for SEM. The images of cell morphology were taken through SEM at 10,000x magnification. Scale bar 2 µm. b) Cell length (µm) for both SecA1 and TatAC was measured, quantified using ImageJ software and plotted using GraphPad Prism software. The statistical analysis was performed by One way ANOVA, **** p<0.0001. c) TEM images of *Rv-vc_dc_* +ATc*, Rv-secA1_dc_* -/+ATc*, Rv-secA1_dc_::secA1* +ATc*, Rv-tatA_dc_* -/+ATc*, and Rv-tatA_dc_::tatAC* +ATc*. Rv-vc_dc_* +ATc*, Rv-secA1_dc_* -/+ATc*, Rv-secA1_dc_::secA1* +ATc*, Rv-tatA_dc_* -/+ATc*, and Rv-tatA_dc_::tatAC* +ATc were grown for 4 days post ATc addition, cells were fixed and processed for TEM. The images of cell membrane were taken at 45,000x magnification. Scale bar 200 nm. d) *Rv-vc_dc_* +ATc*, Rv-secA1_dc_* -/+ATc*, Rv-secA1_dc_::secA1* +ATc*, Rv-tatA_dc_* -/+ATc*, and Rv-tatA_dc_::tatAC* +ATc were grown and cells were harvested for the assays on day 2 and day 4 post ATc treatment. A_600_ ∼0.5 cells were treated with 5 µg/ml PI (Propidium Iodide), incubated for 30 min in dark and washed. The cells were fixed using 4% paraformaldehyde and PI positive cell populations were measured using flow cytometry. Excitation peak at 535 nm and emission peak at 617 nm. The shift in the fluorescence reading is shown here.

To investigate the changes in the permeability of the membrane, we performed Ethidium Bromide (EtBr) and Propidium Iodide (PI) uptake experiments. If the cells have indeed become more permeable after SecA1 and TatAC depletion, we would expect increased EtBr uptake and a shift in the PI fluorescence intensity compared with *Rv-vcdc* +ATC. In line with this hypothesis, we observed increased intensity of EtBr in the cells upon depletion of SecA1 or TatAC for 4 days compared with the control and the 2 days depleted samples (Fig. S6c & d). The increased uptake of EtBr four days post depletion in SecA1 and TatAC depleted samples could be restored to *Rv* levels upon complementation (Fig. S6c & d). For the PI uptake assay, the samples were sorted for the population containing the PI signal using a FACS machine. The results obtained were analogous to those observed in the EtBr uptake assay, with both SecA1 and TatAC-depleted samples showing a shift in fluorescence compared to the undepleted, control, and complemented samples (Fig. 7d). Taken together, depletion of SecA1 or TatAC causes changes in the *Mtb* membrane ultrastructure, which in turn leads to increased permeability.

## Discussion

The manual reconstruction of secretion systems in *Mtb*, encompassing 92 secretory components and 198 mechanistic reactions, organizes diverse information on molecular components, which are often scattered as experimental evidence, and the associated mechanistic interactions involved in the secretion process into a systems-level model. The first genome-scale model for protein secretion was developed for yeast, *Saccharomyces cerevisiae,* using a bottom-up approach with the Protein Specific Information Matrix (PSIM) framework, providing a systems-level understanding of secretion machinery in eukaryotes^52^. Subsequent studies have performed a genome-scale analysis of protein secretion system in *Aspergillus oryzae*^53^ and humans^54^. Notably, a later study^55^ has integrated the core secretory pathway into genome-scale metabolic models for humans (RECON2.2s), mice (iMM1685s), and CHO cells (iCHO2048s) to determine the energetic costs and machinery demands for secreted proteins. In contrast, most available genome-scale models for prokaryotes mainly focus on metabolic pathways and lack detailed representations of protein secretion machineries and their regulation^56–58^. *Mtb* has been extensively studied with models such as iNJ661^59^, GSMN-TB^60^, iEK1011^61^, and sMtb2018^62^, which combine genomic and biochemical data to simulate *Mtb* metabolism under various conditions. However, the available metabolic models for *Mtb* have not been extended to build an integrative model with detailed mechanistic description of protein secretion machinery at the systems-level.

*Mtb* relies on Sec, Tat, and multiple specialized secretion systems, such as the Type VII secretion system (T7SS), which includes five ESX systems (ESX-1 to ESX-5) and SecA2. While reports offer detailed mechanistic insights, the modular organization of T7SS^64^ and the characterization of SecA2^21^ are available independently; however, a single, integrated reconstruction of *Mtb* secretion pathways has not been established. In this study, we aimed to identify all components of secretory systems and to delineate the roles of specific elements (such as SecA1 and Tat) in *Mtb* virulence. Fig. 1 offers a mechanistic characterization of common components in secretion systems, considering the functional evidence from existing experimental data. Instead of relying on computational simulations, we provided a pictorial and conceptual reconstruction that helps visualise the elements of the secretory machinery and their architecture (Fig. 2). The Sec pathway driven by SRP, SecA1 ATPase, and Tat pathway is depicted in our model in detail (Fig. 2 and Fig. S1). By integrating transcriptomic and ChIP-Seq datasets, we identified key transcription factors that regulate the expression of secretion system genes (Fig. 3b), revealing the dynamic regulatory networks that underpin their function, which was absent in most previous studies^35,64,65^. In the future, our reconstruction of *Mtb* secretion systems can be expanded to create a proteome-constrained, genome-scale protein secretory model for *Mtb*, similar to the recently developed model for yeast^66^.

CRISPRi-mediated knockdown can affect genes downstream of the target gene in an operon. While *secA1* is a single-gene operon, the *tatC* gene is downstream of *tatA* gene. Our qRT-PCR experiments showed that the expression of both *tatA* and *tatC* was downregulated upon induction of guide RNA and dCas9 (Fig. 4). Since both TatA and TatC function in the same secretion pathway, we did not attempt to create a specific TatA depletion strain by complementing with *tatC* at another integrative locus. Multiple groups, including ours, have performed mass spectrometric analysis of the mycobacterial secretome. The number of secretory proteins identified in several major studies ranged from 257^47^, 254^48^, 481^50^ to 932 in the current study. The increased numbers in the current study may be due to technical improvements in the mass spectrometer’s sensitivity. Here, we focused on 439 proteins that were common between the current study and one of the three previous studies (Fig. 5). A prior study in *Mycobacterium marinum* and *M. marinum esx-5* null identified 30 proteins, including 24 PE/PPE proteins that depended on the presence of ESX-5^67^. Another study identified a total of 66 proteins (33 increased and 33 decreased) that were either became more or less abundant in the *ΔsecA2* mutant compared with the wild type^22^. Here, we report for the first time that secretion of 118 proteins was dependent on SecA1 and 138 on TatAC systems, respectively (Fig. 5). Interestingly, 72 proteins were commonly impacted by both SecA1 and TatAC depletion (Fig. S4), suggesting either dependence on both pathways due to cross-talk or depletion-dependent perturbations in the cell.

Upon SecA1 depletion, lipoproteins, which are typically localized to the membrane and contain classical Sec signal peptides, were found to be reduced in abundance (Fig. 5). We also identified RipA, PG endopeptidase^68^, and Ami1, an amidase^69^ necessary for maintaining PG synthesis, as being lower in abundance (Fig. 5). In *Streptomyces coelicolor*, the authors identified 43 Tat pathway substrates, 25 of which were validated using a sensitive agarase reporter assay^70^. *Mtb* utilizes the Tat pathway to export various proteins, including enzymes, hydrolases, and proteins involved in cell wall processes^17^. Tat-deficient mutants in *Msm* failed to export β-lactamases such as BlaS to the cell envelope, resulting in sensitivity to β-lactam antibiotics^71^. Rv2525c also plays a role in maintaining cell wall integrity and stress adaptation in *Mtb*^72^. We observed lower abundances of Rv2525c and plcB proteins in the Tat-depleted secretome.

A total of 528^73^, 1417^74^, 1318^22^, and 1539^51^ membrane proteins were obtained from membrane proteome studies conducted by various groups previously, and 2335 were identified in our current study. The variation in the number of membrane proteins may be due to the type of membrane fractionation methods and biological conditions used. Comparing our study with the previous two studies^22,51^ resulted in 943 proteins that were common to our study and one of the other two studies. Among these, 260 were differentially abundant in SecA1, and 81 for TatAC depletion (Fig. 6). A previous study showed that 13 out of 15 solute binding proteins (SBPs), 12 out of 21 Mce transporter family members were reduced, and 13 out of 21 identified DosR-regulated proteins were increased in the *ΔsecA2* mutant^22^. Here, we observed that, upon SecA1 depletion, metabolic proteins, mostly those involved in lipid degradation, were highly abundant, while cell wall synthesis proteins were less abundant (Fig. 6). A similar trend was observed in TatAC depletion. A previous study validated the localization of LpqH, a membrane-anchored lipoprotein^75^, which was lower in abundance in the membrane fraction upon SecA1 depletion in our study (Fig. S5c).

A notable point was that we observed many cytosolic proteins in the secretome upon SecA1 or TatAC depletion, which were not analyzed further because they were not part of the secretome in earlier studies (Fig. S4d & e). We have observed a similar trend even in the membrane proteome upon SecA1 and TatAC depletion (Fig. S5c & d). This observation led to an examination of the ultra-structures by SEM and TEM (Fig. 7). Data Fig. 5-7 suggests that the loss of SecA1 and TatAC appears to disrupt cellular homeostasis, which in turn seems impact the membrane integrity and become more permeable, leading to leakage of periplasmic and cytoplasmic proteins (Fig. 7). Although Sec and Tat pathways are mechanistically distinct, a few proteins can be targeted by either pathway, showing the flexibility of their target signal peptides^6^. Crosstalk is evident in quality control and regulatory processes, where overproduction or dysfunction in any part of the pathway can affect others^16^. This interplay ensures bacteria balance the secretion systems for various proteins crucial for virulence, adaptation, and cellular physiology.

## Materials and Methods

### Bacterial strains and plasmids used for the study

*Mtb H37Rv* (*Rv*), a pathogenic laboratory strain sensitive to all antibiotics, was used in this study. Conditional knockdown strains *Rv-secA1_dc_, Rv-tatA_dc,_* and complementation strains were generated by electroporating relevant constructs into *Rv*. The pTetIntdCas9 plasmid used in this study has dCas9 under tetracycline (ATc) inducible promoter, L5 integrative site, which was described previously^44^. Two guide RNAs each were designed to achieve sufficient depletion upon the addition of ATc. *sec1144* and *sec1183* are the guide RNA sequences aimed at targeting *secA1,* and *tat1142* and *tat1181* are two guide RNA sequences for *tatA*. Two guide RNAs, each of 20 nucleotides complementary sequence for each target gene from 5’UTR and after the PAM sequence 5’NGG3’ were cloned into the pTetIntdCas9 vector pTet-dC-SecA1 and pTet-dC-TatA (Fig. S3a & b). The expression of dCas9 and the target sequence-specific guide RNAs, both under Tet inducible promoter, halts the RNAP from proceeding with transcription; hence, the target gene’s knockdown occurs (Fig. S3c). For complementation, the *secA1* or *tatA* (*tatA_gmut_*) were mutated (Fig. S3d) such that the guide RNA binding is abrogated. *secA1_gmut_* and its native promoter were amplified separately, digested and cloned into pSNG-S vector containing giles attP and Integrase (unpublished) to generate pSNG-secA1_gmut._ *tatA_gmut_*along with downstream *tatC* and their native promoter, were amplified separately (Fig. S3d), digested and cloned into pSNG-S vector to generate pSNG-tatAC_gmut_.

### Lysate preparation and Western blots

*Rv-vc_dc_, Rv-secA1_dc_, Rv-tatA_dc_* strains were grown until A_600_ ∼0.8, and whole cell lysate (WCL) was prepared by resuspending the cell pellet in lysis buffer (1x PBSG, 1x PMSF, and PIC: Protease Inhibitor Cocktail). The cells were lysed using bead beating (8-10 cycles, 1 min ON and 2 min OFF). The cells were centrifuged at 13000 rpm for 10 min at 4°C, and the supernatant was estimated using the BCA reagent (Thermo-Fischer). 40 µg of the WCLs were resolved on 10% (for SecA1) or 15% (for TatA) SDS-PAGE gels, transferred to nitrocellulose membranes, and probed with α-SecA1 or α-TatA, or α-GroEL1 antibodies raised in the lab.

### RNA isolation and qRT-PCR

The *Rv-vc_dc_, Rv-secA1_dc_, and Rv-tatA_dc_* strains, with or without ATc, were grown until A_600_ ∼0.8, and cells equivalent to 10 O.D. were harvested after 3 days of ATc treatment. The cell pellets were resuspended in 1 ml of Trizol (Invitrogen), and cells were lysed by bead beating (3 rounds of 30 sec ON and 2 min OFF). The cells were centrifuged at 13000 rpm for 10 min at 4°C, and 300 µl of chloroform was added to the supernatants. The samples were vortexed, centrifuged, and the aqueous layer containing RNA was precipitated. The pellet was resuspended in RNase-free water, treated with DNase I (Invitrogen), and purified using a Qiagen column. 1 µg of RNA was reverse transcribed using the iScript cDNA synthesis kit (Bio-Rad). Quantitative real-time PCR was performed with iTaq Universal SYBR Green Supermix (Bio-Rad) on the QuantStudio 3 system (Applied Biosystems). Data were normalized to 16S rRNA expression levels. The ΔΔCt algorithm was used to evaluate the relative fold change in gene expression.

### Bacterial survival *in vitro* and upon *ex vivo* peritoneal macrophage infection

The strains were inoculated at A_600_ ∼0.1 in the presence and absence of ATc. The CFUs were enumerated on day 0 (data not shown) and day 4. Colonies were calculated and plotted using GraphPad Prism. For the *ex vivo* infections, peritoneal macrophages were used. Briefly, thioglycolate was injected into the mouse peritoneum, and macrophages were isolated 4 days post-injection (Fig. 4d). 3 x 10^5^ cells were seeded per well, and infection was performed at a 1:5 (macrophage:bacterial cells) MOI. CFUs were enumerated at 4 and 72 h post-infection with and without ATc. Colonies were counted, and a graph was plotted using GraphPad Prism.

### Reconstruction of Secretory systems in *Mtb*

Inspired by the work by Feizi *et al*. on genome-scale reconstruction of protein secretion system in yeast *Saccharomyces cerevisiae*, we created a manually curated reconstruction of the secretion systems in *Mtb*^52^. Note that yeast is a eukaryote while *Mtb* is a bacterium. To this end, a systematic manual curation of existing literature on bacterial secretion systems, supported by both experimental and/or computational evidence, was performed as depicted in the workflow^52,76^ (Fig. 1a). Based on the compiled evidence, components of secretion pathways namely, Sec dependent (SRP dependent, SecA1 dependent, and SecA2 dependent),

Sec independent (Tat), and T7SS were collated and annotated (Table S1 and S2). Further, the protein signatures relevant for secretion and localization, such as presence or absence of signal peptide sequences, lipo motif (LM), transmembrane domain (TMD), etc. were assigned using the computational tools SignalP v6.0^77^, Phobius v10.1 (https://phobius.sbc.su.se/), DeepTMHMM v1.0 (https://services.healthtech.dtu.dk/services/DeepTMHMM-1.0/) and PcnsHub^78^, facilitating the identification and classification of secreted proteins (Fig. 1a). Detailed mechanistic information from the existing literature was used to further organize the secretion process into a series of pseudo-chemical reactions, similar to reactions formulated by Feizi *et al.* for classical secretion pathway in Yeast^52^. A pictorial representation of these reactions was created using Biorender (Fig. 2, Fig. S1 & S2).

To explore the transcriptional regulatory control of secretory components, the publicly available ChIP-Seq data^40^ for *Mtb* transcription factors (TFs) was used to determine the TFs directly controlling the different *Mtb* secretory components (Fig. 3, Table S3). A visualization of TFs controlling the *Mtb* secretory components was generated using the R package ggplot2 (https://ggplot2.tidyverse.org/). The gene-level expression changes of the secretory components under different stress conditions were obtained from a transcriptome study performed by Khan et al., 2024^42^. The differential expression was viewed using a heatmap constructed using R package, pheatmap (https://cran.r-project.org/web/packages/pheatmap/index.html).

### Culture filtrate protein preparation

To isolate the culture filtrate (CF) proteins from *Mtb*, *Rv-vc_dc_, Rv-secA1_dc_,* and *Rv-tatA_dc_*were grown till early log-phase culture with and without ATc (500 ng/mL) in Sauton’s media in quadruplets^45^. Briefly, the cells were harvested and centrifuged at 4000 rpm for 10 min at 4°C. The supernatants were collected into a fresh 50 mL falcon, and pellets were kept aside for preparation of WCL (WCL: Whole Cell Lysate). The supernatants were centrifuged again following the same condition and repeated twice. The supernatants were collected and passed through a 0.2 µm syringe filter at the last centrifuge. The filtered supernatants (CF proteins) were concentrated using 3 kDa Amicon centricons (∼150 mL CF to 1 mL) and stored at -80°C.

### Secretome sample preparation for Mass spectrometry

For mass spectrometry of CF proteins, 100 µg were taken into fresh low protein binding MCTs (microcentrifuge tubes), and 6X volume of ice-cold acetone was added for overnight precipitation at -20°C. The samples were centrifuged at 13000 rpm for 10 min at 4°C, supernatant was discarded, and the pellets were dried in a speed vacuum. 100 µl resuspension buffer containing 8M urea and 25 mM ammonium bicarbonate was added to the dry pellet, and the sample was resuspended by vortexing for 30 min at RT. 10 mM of TCEP was added to the tubes and incubated at RT for 30 min. 25 mM of IAA was added to the tubes and incubated at RT for 30 min at dark. The proteins were then subjected to trypsin digestion (Trypsin: protein - 1:20) at 37°C overnight. The peptides were desalted using C-18, eluted in a fresh MCT, and stored at -80°C.

### Protocol for isolation of membrane and cytosolic fractions

*Rv-vc_dc_, Rv-secA1_dc_,* and *Rv-tatA_dc_* were grown in the presence and absence of ATc (500 ng/mL) in 7H9+OADC media in quadruplets. On the fourth day, the cells were pelleted and resuspended in lysis buffer (1x PBSG, 1x PMSF, and PIC: Protease Inhibitor Cocktail), and cell lysates were prepared using bead beating for eight rounds (1 min ON and 2 min OFF cycles on ice). Samples were centrifuged at 13000 rpm for 10 min at 4°C. The supernatant was collected and subjected to ultracentrifugation at 100000X g for two h at 4°C. After the first round of ultracentrifugation, the supernatant was collected and designated as the Cytosolic fraction (C). The pellets were resuspended in 1x PBSG with 1X PMSF and subjected to another two rounds of ultracentrifugation to collect the membrane fractions (M). After the final (third) round of ultracentrifugation, the pellets (M) were resuspended in resuspension buffer (5% SDS, 0.1M Tris-Cl pH-8.5) and stored at -80°C.

### MS Sample preparation and processing

25 µg of proteins from the membrane and cytosolic fractions were reduced and alkylated using 5 mM TCEP and 50 mM iodoacetamide separately. Trypsin was then added at a ratio of 1:50 in µg to lysate and incubated at 37°C for 16 h. A C-18 silica cartridge was used to purify the digested peptide, and a speed vacuum was used to dry it. Buffer A (2% acetonitrile + 0.1% formic acid) was used to suspend the dry pellet and submitted to a mass spectrometer containing an Easy-nLC-1000 system connected to an Orbitrap Exploris. The C-18 column (15 cm/1.9 μm) was loaded with 1 µg peptide and was later separated as a gradient using buffer B (80% acetonitrile + 0.1% formic acid) and a flow rate of 500 nL/min. The duration of the LC gradients was 110 min. The eluate was injected for MS analysis. The MS1 spectra were obtained (Max IT = 60ms, AGQ goal = 300%; RF Lens = 70%; R=60K, mass range = 375−1500; profile data). All charge states for a certain precursor were excluded for 30 sec using dynamic exclusion. The top 20 peptides’ MS2 spectra were gathered using the following parameters: AGC target 100%, R = 15K, and Max IT = 60ms.

### Mass spectrometry data analysis and visualization

RAW files were annotated using Proteome Discoverer 2.5 with UniProt database^79^. The precursor and fragment mass limitations of 10 ppm and 0.02 Da were set for dual Sequest and Amanda searches. The trypsin/P enzyme specificity of the protease was set for cleavage at the C terminus of “K/R: unless followed by “P”. The protein false discovery rate (FDR) and peptide spectrum match (PSM) were set at 0.01. Further analysis was carried out by mapping the peptide sequence obtained in culture filtrate and membrane fraction to the respective UniProt ID, and LFQ values were obtained from MaxQuant^80^.

Mycobrowser^30^ database was used for creating a list of *Mtb* proteins with their respective UniProt ID, Locus ID and functional classification, and was augmented with information on the presence or absence of signal peptide, transmembrane helices and non-classical secretion obtained by *in silico* prediction using the protein sequences from Mycobrowser and computational tools SignalP v 6.0^77^, DeepTMHMM v 1.0 (https://services.healthtech.dtu.dk/services/DeepTMHMM-1.0/), Phobius v 10.1 (https://phobius.sbc.su.se/) and PncsHub^78^. The UniProt IDs and the LFQ values obtained from MS data were mapped with the list. The proteins having ≤2 unique peptides were removed. In a similar fashion, the membrane proteome data was mapped with the created list. After filtration, the proteins and LFQ values were log normalized and the differential abundance was performed using limma^81^ and significant proteins were considered for further analysis.

The differential abundance, signal peptide distribution and functional annotation were visualized using pheatmap (https://cran.r-project.org/web/packages/pheatmap/index.html), while volcano plots, alluvial plots and chord plots were generated using ggplot2 (https://ggplot2.tidyverse.org/), ggalluvial^82^, and circlize^83^packages respectively in R.

### Scanning Electron Microscopy (SEM)

For SEM, 5 mL cells at A_600_ ∼0.5 of *Rv-vc_dc_, Rv-secA1_dc_, Rv-secA1_dc_:: secA1, Rv-tatA_dc_,* and *Rv-tatA_dc_::tatAC* were centrifuged at 4000 rpm for 10 min at RT. The supernatant was discarded, and the pellets were resuspended in fixative. The cells were incubated at 37°C. The cells were centrifuged for 10 min, and the pellets were washed twice with 0.1 M Na-Cacodylate buffer. The cells were resuspended in 1% OsO_4_ (Osmium tetroxide) and incubated at RT for 90 min in a twirler. The cells were centrifuged again for 10 min at RT. The supernatant was discarded, and dehydration was performed gradually with different concentrations of ethanol: 25% ethanol for 5 min, 50% ethanol for 7 min, 75% ethanol for 10 min, 95% ethanol for 20 min, and 100% ethanol for 30 min, three times. After each step, samples were centrifuged for 10 min at RT. The pellets were resuspended in HMDS (Hexamethyldisilazane). The samples were subjected to gold coating followed by SEM (S3400N, Hitachi) imaging. Cell lengths were calculated using ImageJ software, and the data were plotted using GraphPad Prism.

### Transmission Electron Microscopy (TEM)

For TEM sample preparation, the SEM protocol mentioned above was followed until the dehydration step. After that, the pellets were resuspended in propylene oxide and incubated at 4°C for 1-2 h on a rotator. The samples were spun down, and the supernatant was discarded. A mixture of propylene oxide and resin in a 1:1 ratio was added to the samples, which were then incubated at 4°C for 2 h in a rotator. This step was repeated with 1:2 and 1:3 propylene oxide to resin ratios, respectively. Finally, pure resin was added to the samples and incubated overnight at 4°C on a rotator. The next day, the resin was discarded, and pure resin was added to the samples, which were incubated at 60°C for 2 h. The samples were then subjected to microtome for transverse sectioning at 70 nm thickness. The sections were stained with uranyl acetate, and images were taken using TEM (Talos L120c, Thermo Fischer Scientific).

### Ethidium bromide (EtBr) uptake assay

*Rv-vc_dc_, Rv-secA1_dc_, Rv-secA1_dc_::secA1, Rv-tatA_dc_,* and *Rv-tatA_dc_::tatAC* strains were inoculated at A_600_ ∼0.1 with and without ATc. The cells were then harvested on day 2 and day 4 for the EtBr uptake assay. The cell pellets were washed with 1 X PBST_80_. Cells at A_600_ ∼0.5 were drawn from the cultures in an MCT, and 1 µg/ml of EtBr was added to the cultures. The cultures were incubated at 37°C for 15 min (data not shown) and 30 min in a standing incubator in the dark. After incubation, the cells were washed with 1 x PBST_80_ twice to remove the residual EtBr. After the second wash, the pellets were resuspended in 4% paraformaldehyde (PFA) for fixation. The cell suspensions were transferred into a 96-well plate to take the fluorescence reading. The cell suspensions were excited at 510 nm, and the emission spectra were collected at 595 nm. Without EtBr, added cells were taken as a control to normalize and get the absolute unit (A.U.) value using GraphPad Prism software.

### Propidium Iodide (PI) uptake assay

*Rv-vc_dc_, Rv-secA1_dc_, Rv-secA1_dc_::secA1, Rv-tatA_dc_, and Rv-tatA_dc_::tatAC* strains were grown in the presence and absence of ATc. On Day 2 and Day 4, the cells were processed for a PI uptake assay. The cells were washed with 1 X PBST_80_. A volume of 5 µg/ml of PI was added to cells at A_600_ ∼0.5 and incubated at 37°C for 5 min in a standing incubator in the dark. After incubation, the cells were washed twice with 1X PBST_80_ to remove any residual PI. The cells were then resuspended in 4% paraformaldehyde (PFA). The cell suspension was used for flow cytometry analysis. The fluorescence of the PI-containing cells was plotted using GraphPad Prism.

## Resource Availability

### Lead contact

Reagent inquiries will be directed to, and fulfilled by, the lead contact, Dr. Vinay Kumar Nandicoori.

### Materials availability

The materials generated in this study are available from the lead contact with a completed materials transfer agreement.

### Data and code availability

The mass spectrometry data generated in this study have been deposited to PRIDE: PXD066777.

Any additional information required to reanalyse the data reported in this paper is available from the lead contact upon request.

## Acknowledgement

Research in this publication was supported by JC Bose Award (JCB/2019/000015), Govt. of India to VKN. VKN and JU acknowledges the support from Department of Biotechnology (DBT) (Grant No. BT/PR50983/MED/29/1650/2023). VKN, PS and YB acknowledge the support from National Institute of Health (NIH) (Grant No. 7R01A|142672-05). AS acknowledges funding from the Department of Atomic Energy (DAE), Government of India, via Apex project to The Institute of Mathematical Sciences (IMSc), Chennai. PS is INSPIRE Fellow from Department of Science & Technology (DST), Government of India (DST/INSPIRE Fellowship/2018/F180904). We are grateful to A. Harikrishna, CSIR-Centre for Cellular and Molecular Biology (CCMB) for his assistance in microtome and TEM image acquisition. We thank CCMB central research facilities: DNA Sequencing, BSL-3, Tissue Culture, Animal House (IAEC-25/2023) and Electron Microscopy, for providing access. Portions of the abstract and cover letter were refined for style and grammar with the assistance of ChatGPT (OpenAI; August 2025). The authors reviewed and edited all text and accept full responsibility for the content. No AI tools were used to generate or alter data, analyses, results, or figures/images.

---

**Supplementary Figure S1: Schematic Representation of protein export through SecA1 dependent Sec pathway and Tat pathway.**

Pictorial model depicting the steps involved in SecA1 dependent secretion and TAT pathway when a protein contains: (a,d) N-terminal Sec/RR-motif signal sequence, Lipo motif and Transmembrane domain, (b,e) N-terminal Sec/RR-motif signal sequence and Lipo motif. (c,f) N-terminal Sec/RR-motif signal sequence and Transmembrane domain. The steps shown were based on the mechanistic reactions framed using the experimental evidence provided in Table S2. Created in BioRender. Nandicoori, V. (2025) https://BioRender.com/fy6pdxr.

**Supplementary Figure S2: Schematic Representation of protein export through SRP dependent Sec pathway.**

Pictorial model depicting the steps involved in SRP dependent secretion when a protein contains: (a) N-terminal SRP signal sequence, Lipo motif and Transmembrane domain, (b) N-terminal SRP signal sequence and Lipo motif, (c) N-terminal SRP signal sequence and Transmembrane domain, and (d) only N-terminal SRP signal sequence. The steps shown were based on the mechanistic reactions framed using the experimental evidence provided in Table S2. Created in BioRender. Nandicoori, V. (2025) https://BioRender.com/6e4dlep.

**Supplementary Figure S3: CRISPRi mediated depletion strains of SecA1 and TatAC.**

a) The vector maps represent here are of pTetInt-dCas9 vector for both SecA1 and TatA. b) The image shows the length of *secA1* (2850 bp) and *tatA* (252 bp) having the PAM sequences and guide RNAs binding regions. The yellow bars represent the PAM sequences and the red bars represent the respective guide RNA s for SecA1 and TatA. c) Schematic representation of CRISPRi mediated transcription inhibition upon addition of ATc. Created in BioRender. Nandicoori, V. (2025) https://BioRender.com/n0g7fiw. d) Diagram depicts the mechanism of complementation strains having guide RNA mutations in both *secA1* (*secA1_gmut-1/2_*) and *tatA* (*tatA_gmut1/2_*). e) Whole cell lysates (WCLs) were prepared from *Rv-vc_dc_*+ATc, *Rv-secA1_dc_*-/+ATc, *Rv-secA1_dc_::secA1* +ATc, *Rv-tatA_dc_*-/+ATc and *Rv-tatA_dc_::tatAC* +ATc from grown 4^th^ day culture. 50 µg of each protein lysates were resolved on 10% (SecA1) 15% (TatA) SDS-PAGE, transferred on Nitrocellulose membrane and probed with α-SecA1 (for SecA1 depletion), and α-TatA (for TatA depletion) antibodies generated in the lab. α-GroEL1 antibody was used as loading control.

**Supplementary Figure S4: Isolation and validation of CF proteins from SecA1 and TatAC.**

a) Schematic diagram of CF protein isolation from *Mtb Rv-vc_dc_, Rv-secA1_dc_,* and *Rv-tatA_dc_* cells in the presence of ATc. Created in BioRender. Nandicoori, V. (2025) https://BioRender.com/fauewmb. b) 50 µg of the isolated CFs fractions and WCL were resolved on SDS-PAGE, proteins were transferred onto Nitrocellulose membrane and probed with α-PknB (cytosolic protein), α-GroEL1 (cytosolic protein) and α-Ag85B (CF protein). Coomassie was performed as protein loading control. c) Venn diagram represents the comparison of our secretome data with four other studies performed by other groups available in the literature. d) The table represent the list of unique proteins identified in the secretome of *Rv-secA1_dc_*+ATc and *Rv-tatA_dc_*+ATc. The proteins highlighted in the green box represent the high abundant proteins and proteins highlighted in the red box represented the low abundant proteins. e) The table represent the list of common proteins identified in the secretome of *Rv-secA1_dc_*+ATc and *Rv-tatA_dc_*+ATc. The proteins highlighted in the green box represent the high abundant proteins and proteins highlighted in the red box represented the low abundant proteins.

**Supplementary Figure S5: Validation of membrane proteome of SecA1 and TatAC membrane fractions.**

a) Schematic representation of isolation of membrane and cytosolic fractions from *Rv-vc_dc_, Rv-secA1_dc_,* and *Rv-tatA_dc_* in presence of ATc. Created in BioRender. Nandicoori, V. (2025) https://BioRender.com/a0uooni. b) 50 µg of the isolated membrane, cytosol and WCL fractions were resolved on SDS-PAGE, proteins were transferred in the Nitrocellulose membrane and probed with α-PknB (membrane protein) and α-RpoA (cytosolic protein). c) The table represent the list of unique proteins identified in the membrane proteome of *Rv-secA1_dc_*+ATc and *Rv-tatA_dc_*+ATc. The proteins highlighted in the green box represent the high abundant proteins and proteins highlighted in the red box represented the low abundant proteins. e) The table represent the list of common proteins identified in the membrane proteome of *Rv-secA1_dc_*+ATc and *Rv-tatA_dc_*+ATc. The proteins highlighted in the green box represent the high abundant proteins and proteins highlighted in the red box represented the low abundant proteins.

**Supplementary Figure S6: Validation of membrane alteration upon SecA1 and TatAC depletion.**

a) The bars represent the % of cells having a cell length of <3 µm, 3-4 µm and >4 µm in *Rv-vc_dc_* +ATc, *Rv-secA1_dc_* -/+ATc, *Rv-secA1_dc_::secA1* +ATc, *Rv-tatA_dc_* -/+ATc and *Rv-tatA_dc_::tatA* +ATc. b) The bars depict the percentage of bulged cells vs normal cells in *Rv-vc_dc_* +ATc, *Rv-secA1_dc_* -/+ATc, *Rv-secA1_dc_::secA1* +ATc. c) The graphs represent the fluorescence (A.U.) upon EtBr uptake by *Rv-vc_dc_* +ATc, *Rv-secA1_dc_* -/+ATc, *Rv-secA1_dc_::secA1* +ATc strains on Day 2 and Day 4. d) The graphs represent the fluorescence (A.U.) upon EtBr uptake by *Rv-vc_dc_* +ATc, *Rv-tatA_dc_* -/+ATc, *Rv-tatA_dc_::tatA* +ATc strains on Day 2 and Day 4.

