## Supplementary figures for "Sec and Tat mediated secretion safeguards *Mycobacterium tuberculosis* membrane homeostasis"

### Supplementary Figure 1.

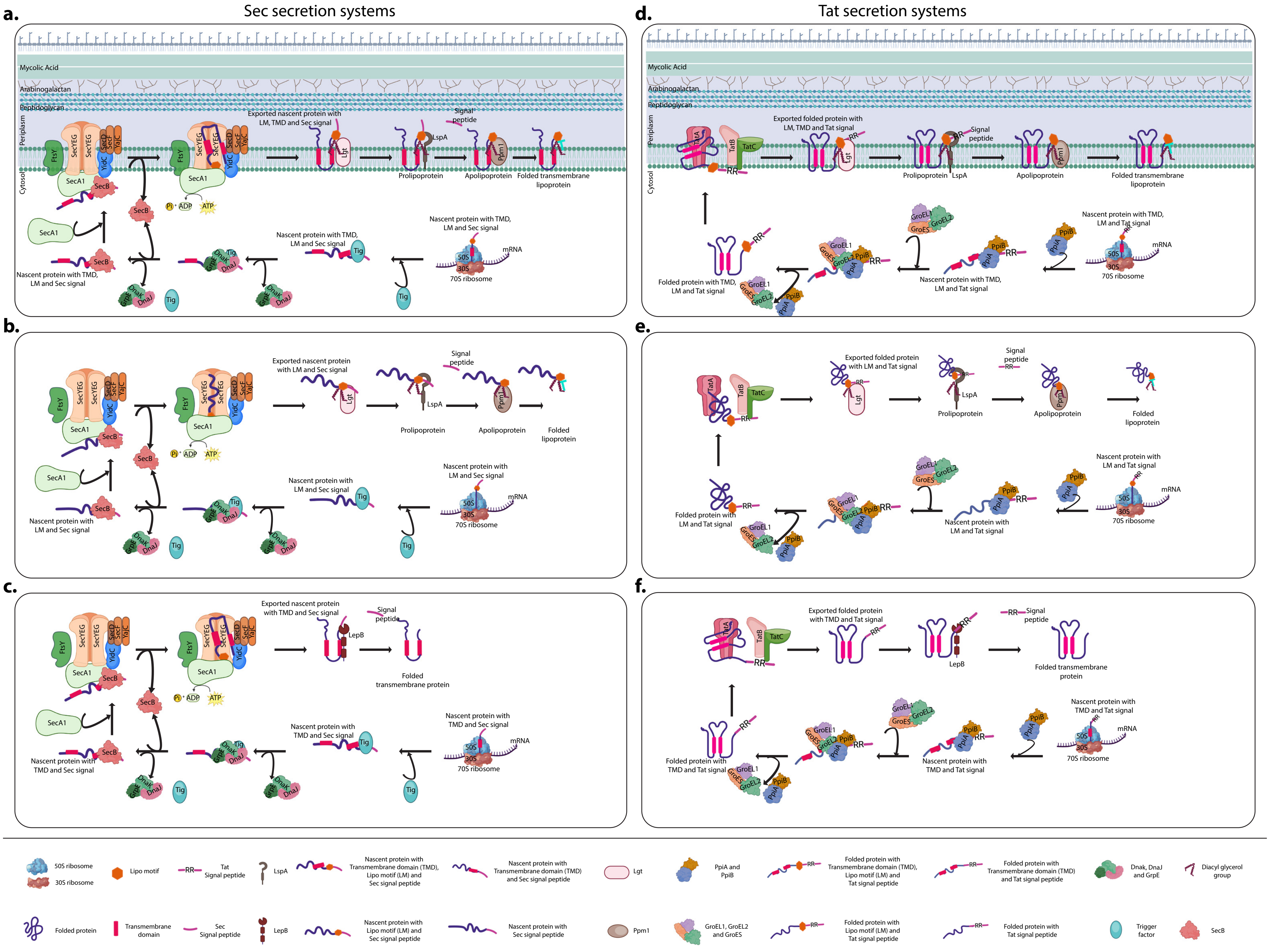

**Supplementary Figure 2.**

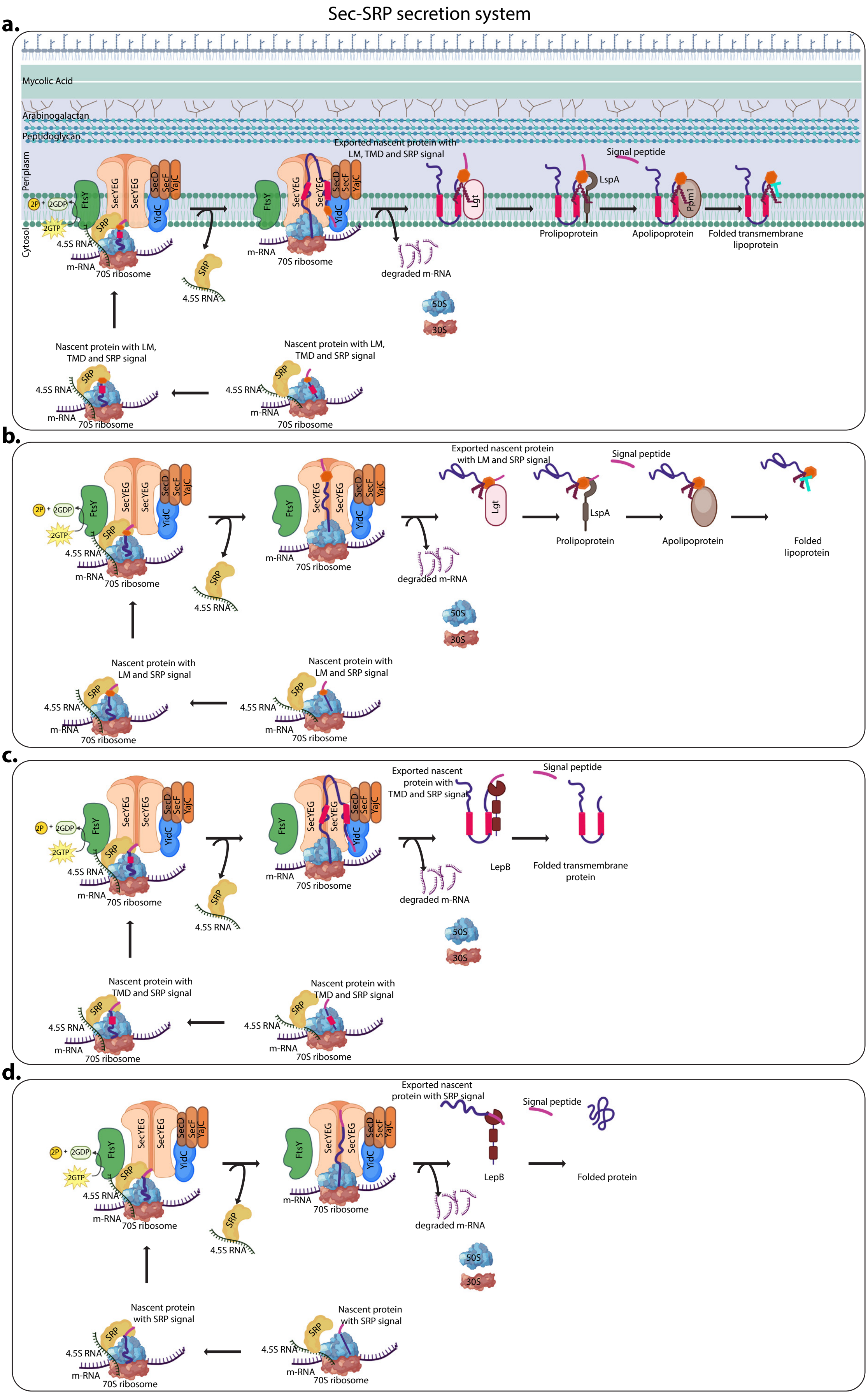

#### Supplementary Figure 3

**a.**

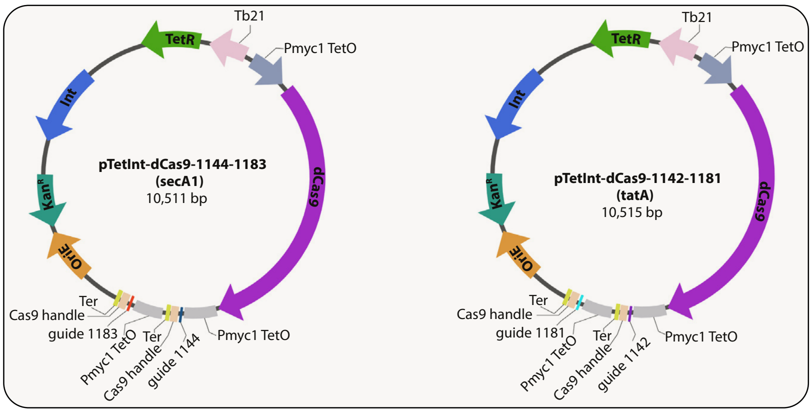


**b.**

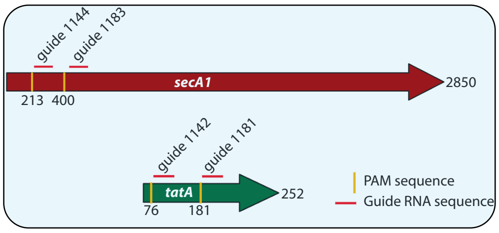


**c.**

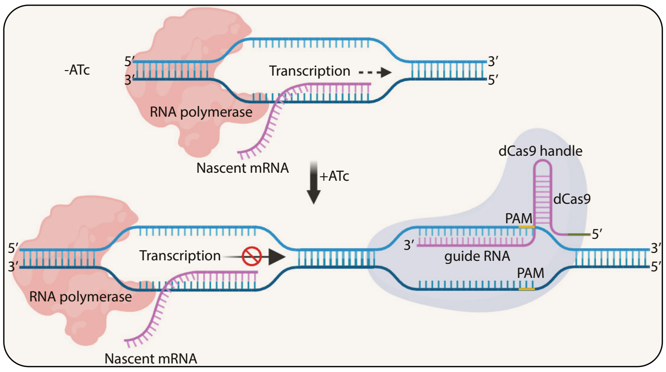


**d.** Mutations for guide RNA complementation

*secA1*$_{WT}$ CCC GAG GCC TTC GCC GTG GCC CGC

*secA1*$_{gmut-1}$ CCC GA**A** GC**T** TT**T** GC**T** GT**A** GC**T** CG**G**

*secA1*$_{WT}$ C CTG GCT AAA CGC GAC AGT GAG T

*secA1*$_{gmut-2}$ **T** CT**C** GC**A** AA**G** CG**G** GA**T** AG**C** GA**A** T

*tatA*$_{WT}$ CC GAT GCG GCG CGC TCG TTG GGC

*tatA*$_{gmut-1}$ CA GA**C** GC**C** GC**C** CG**G** TC**C** TT**A** GG**G**

*tatA*$_{WT}$ CG GTG CAA TCG CAG CGT GTC GAC CCC

*tatA*$_{gmut-2}$ CA GA**C** CA**G** TC**C** CA**A** CG**A** GT**G** GA**T** CC**G**

**e.**

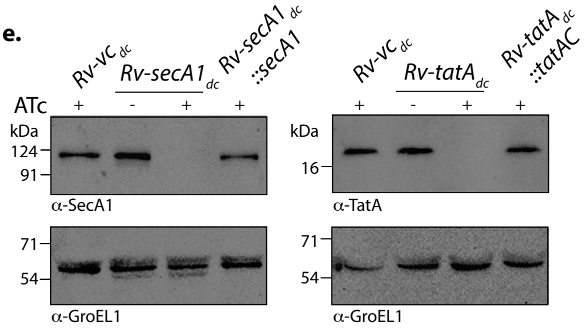

#### Supplementary Figure 4

**a.**

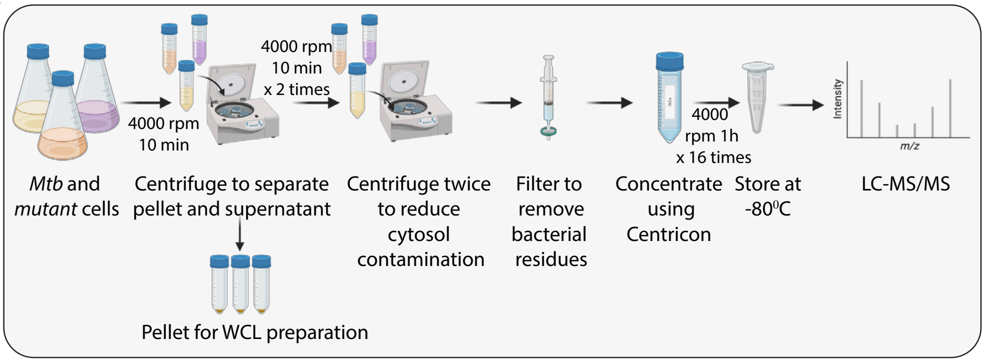


**b.**

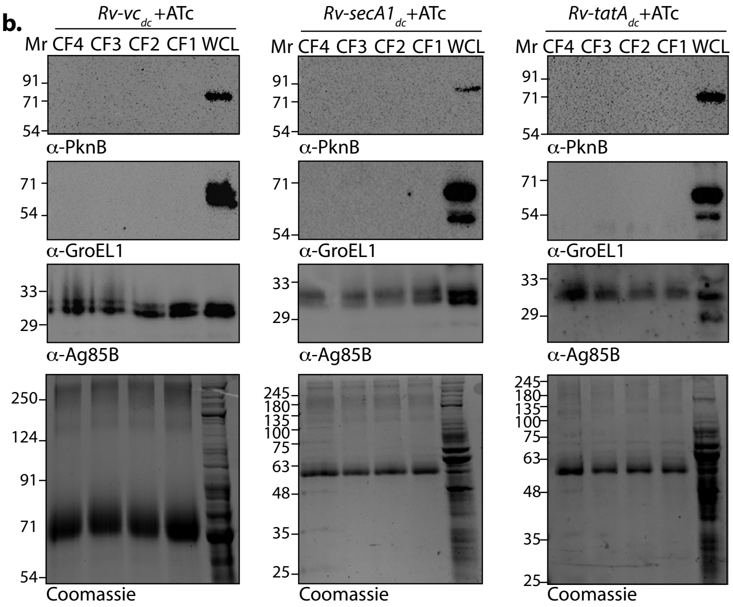


**c.**

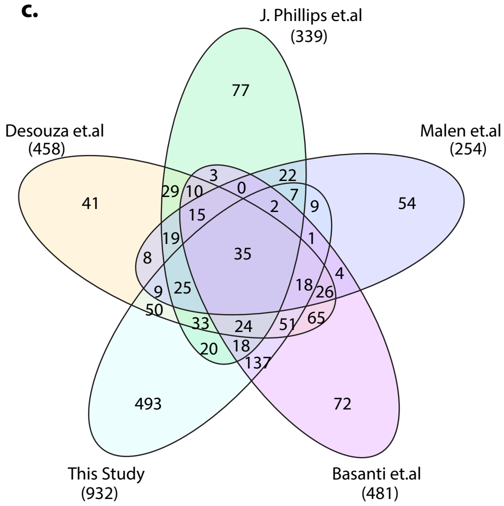


**d.**

| List of unique proteins identified in secretome of depletion strains | | | | | | |
|---|---|---|---|---|---|---|
| *Rv-secA1*$_{dc}$ +ATc | | | *Rv-tatA*$_{dc}$ +ATc | | | |
| *fadD14* | *Rv2161c* | *ripA* | *Rv0566c* | *Rv2466c* | *rmlC* | *ldtA* |
| *Rv2477c* | *ilvE* | *Rv0888* | *nusG* | *Rv2971* | *hadC* | *Rv0680c* |
| *Rv1248c* | *amiC* | *pstS1* | *purH* | *glpK* | *Rv3684* | *Rv0518* |
| *rpoC* | *hisD* | *Rv0526* | *Rv3075c* | *fadE15* | *Rv0333* | *Rv1184c* |
| *Rv0088* | *Rv1462* | *Rv2672* | *nuoG* | *Rv0546c* | *Rv3282* | *lpqD* |
| *Rv0187* | *ndkA* | *lpqE* | *pheT* | *leuA* | *add* | *lprC* |
| *mmaA1* | *ald* | *Rv0309* | *fadA* | *hadB* | *Rv0580c* | *lprE* |
| *ilvC* | *rraA* | *lppO* | *greA* | *gatA* | *pepB* | *Rv0063* |
| *Rv2603c* | *gpm2* | *glnH* | *gcvB* | *Rv2993c* | *Rv3720* | *Rv0774c* |
| *fas* | *fadE35* | | *nuoC* | *dhaA* | *Rv1771* | *Rv2525c* |
| *Rv2302* | *thiC* | | *Rv2676c* | *Rv0183* | *ppsE* | *Rv0846c* |
| *Rv3627c* | *Rv2226* | | *prfB* | *Rv0731c* | *cyp142* | |
| *moxR1* | *sapM* | | *prpD* | *fadA5* | *PPE51* | |
| *tgs2* | *dacB2* | | *Rv2454c* | *espF* | *Rv1910c* | |
| *Rv1543* | *TB22.2* | | *trxB2* | *Rv0390* | *Rv1488* | |
| *Rv2050* | *Rv1987* | | *mtrA* | *katG* | *pepQ* | |
| *purC* | *Rv3572* | | *adk* | *lprQ* | *Rv3400* | |
| *php* | *Rv0455c* | | *trpE* | | *PPE20* | |
| | *Rv2668* | | *guaA* | | *espR* | |

**e.**

| List of common proteins identified in secretome of depletion strains | | | |
|---|---|---|---|
| *adhD* | *purA* | *espC* | *Rv0477* |
| *ahpD* | *rpiB* | *gadB* | *Rv0519c* |
| *atpG* | *Rv0036c* | *glnB* | *Rv0559c* |
| *dapD* | *Rv0146* | *lppW* | *Rv0787* |
| *efp* | *Rv1377c* | *lpqL* | *Rv1061* |
| *fadD28* | *Rv1463* | *lpqR* | *Rv1257c* |
| *fadD31* | *Rv1558* | *lprK* | *Rv1260* |
| *fadE5* | *Rv1794* | *mctB* | *Rv2190c* |
| *fadE6* | *Rv2229c* | *mmpS5* | *Rv2223c* |
| *fixA* | *Rv2260* | *mpt70* | *Rv2253* |
| *frr* | *Rv3688c* | *mpt83* | *Rv3033* |
| *fusA1* | *sigK* | *PE15* | *Rv3395A* |
| *fusA2* | *sodA* | *PE31* | *Rv3491* |
| *gcpE* | *tal* | *PE5* | *Rv3717* |
| *gltS* | *TB16.3* | *plcB* | |
| *icl1* | *tesA* | *ponA2* | |
| *lpdA* | *tig* | *PPE41* | |
| *mas* | *trxC* | *Rv0356c* | |
| *mcr* | *umaA* | *Rv0398c* | |
| *pgk* | | | |

#### Supplementary Figure 5

**a.**

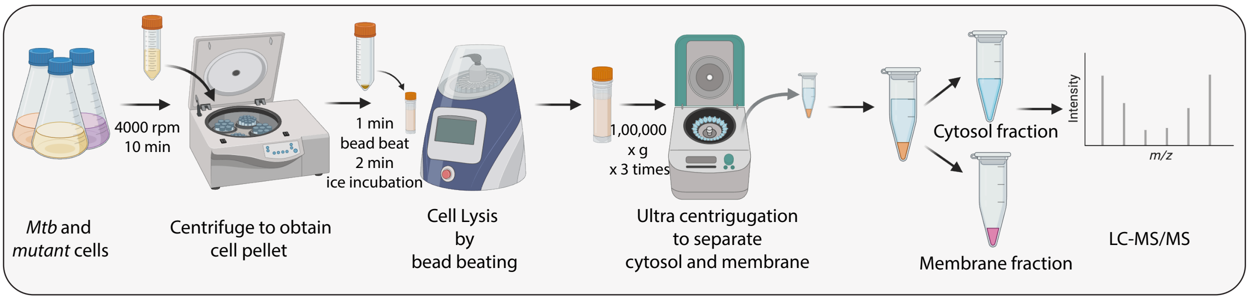


**b.**

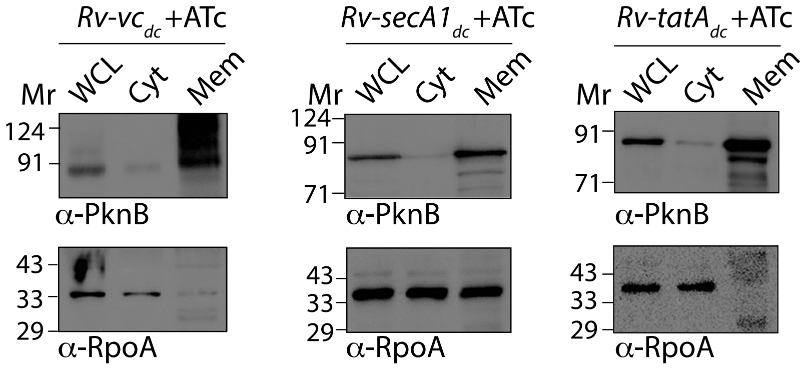


**c.**

| List of unique proteins identified in membrane proteome of depletion strains | | | | | | |
|---|---|---|---|---|---|---|
| *Rv-secA1*$_{dc}$ +ATc | | | | *Rv-tatA*$_{dc}$ +ATc | | |
| *hsp* | *pks13* | *lppJ* | *yrbE1B* | *Rv2799* | *kstR* | *nrdF1* |
| *Rv1566c* | *fbpC* | *Rv0901* | *lpqH* | *Rv1510* | *Rv2958c* | *ripA* |
| *fadD26* | *mas* | *Rv0312* | *Rv2599* | *cyp121* | *Rv2862c* | *Rv3627c* |
| *carB* | *pyrH* | *Rv3673c* | *PE31* | *Rv2264c* | *Rv1993c* | *Rv2361c* |
| *ahpC* | *nusG* | *Rv0696* | *Rv3233c* | *Rv1179c* | *gpsI* | *pepA* |
| *prcA* | *nrdE* | *irtB* | | *Rv1069c* | *sodA* | *Rv2567* |
| *gcvT* | *fadE21* | *Rv1424c* | | *plcB* | *qcrB* | *Rv1730c* |
| *aroF* | *lppD* | *lppK* | | *Rv3226c* | *pks5* | |
| *Rv1875* | *Rv3902c* | *Rv3127* | | *mce4A* | *pitA* | |
| *fadD23* | *Rv0801* | *Rv1272c* | | *vapC27* | *Rv2113* | |
| *Rv2230c* | *nusB* | *espC* | | *citE* | *qcrA* | |
| *fadD13* | *accD1* | *dctA* | | *Rv2219A* | *Rv0158* | |
| *Rv2477c* | *Rv3767c* | *tsnR* | | *espE* | *Rv0738* | |
| *devB* | *mpa* | *Rv0201c* | | *Rv2402* | *Rv3587c* | |
| *ilvC* | *Rv2728c* | *cdh* | | *Rv1100* | *Rv0888* | |
| *adhC* | *Rv3267* | *Rv3725* | | *Rv1456c* | *Rv3277* | |
| *dapD* | *Rv1546* | *yrbE1B* | | | *TB27.3* | |
| *Rv1874* | *folE* | *lpqH* | | | *Rv2844* | |
| *Rv1465* | *Rv0245* | *Rv2599* | | | *ggtA* | |
| *fbpA* | *ndh* | *PE31* | | | *pks6* | |
| *guaB1* | *Rv3684* | *Rv3233c* | | | *qcrC* | |
| *accD2* | *Rv2250c* | | | | *Rv2962c* | |
| *hisC2* | *fadD25* | | | | *mshD* | |
| *Rv0347* | *Rv1869c* | | | | *Rv0877* | |
| *glnA2* | | | | | *stp* | |
| *trpD* | | | | | *Rv2843* | |

**d.**

| List of common proteins identified in membrane proteome of depletion strains | |
|---|---|
| *Rv3096* | *lpqP* |
| *Rv2038c* | *Rv0539* |
| *prcB* | *Rv1828* |
| *Rv2877c* | *Rv3714c* |
| *ppsE* | *Rv3707c* |
| *Rv2553c* | *Rv2102* |
| *Rv2324* | *Rv2820c* |
| *ceoB* | *Rv3076* |
| *Rv0455c* | *rpsT* |
| *Rv1996* | *Rv3238c* |
| *inhA* | *lpqE* |
| *tatA* | *Rv2468c* |
| *Rv2693c* | *lpqK* |
| *Rv2170* | |
| *Rv1896c* | |
| *aroA* | |
| *Rv0145* | |
| *pyrE* | |
| *Rv2455c* | |

#### Supplementary Figure 6

**a.**

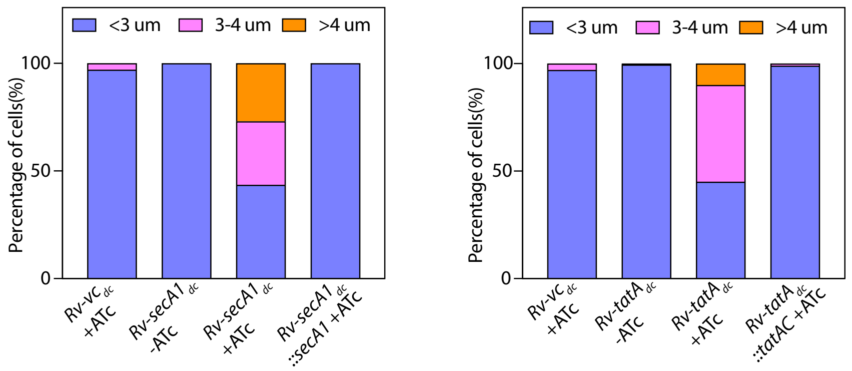


**b.**

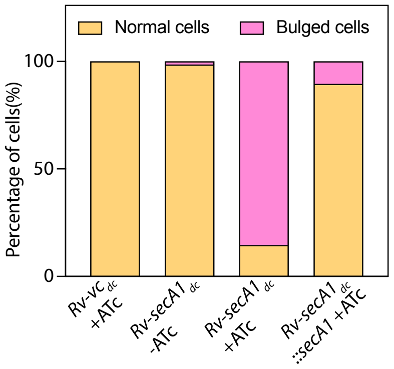


**c.**

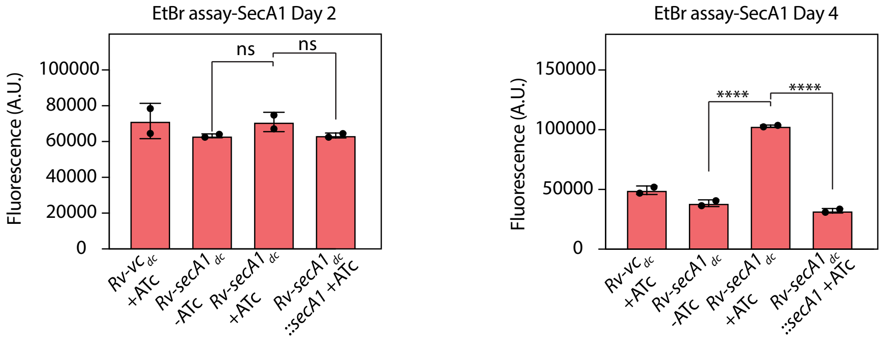


**d.**

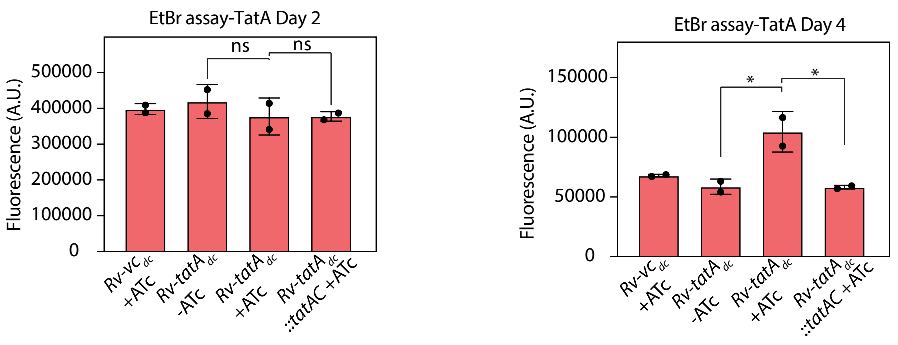
